# Phenylpropionc acid produced by gut microbiota alleviates acetaminophen-induced hepatotoxicity

**DOI:** 10.1101/811984

**Authors:** Sungjoon Cho, Xiaotong Yang, Kyoung-Jae Won, Vanessa Leone, Nathaniel Hubert, Eugene Chang, Eunah Chung, Joo-Seop Park, Grace Guzman, Hyunwoo Lee, Hyunyoung Jeong

**Author notes:** Corresponding authors **Correspondence:** Hyunyoung Jeong, PharmD, PhD, Hyunwoo Lee, PhD, Department of Pharmaceutical Sciences, University of Illinois at Chicago, 900 South Ashland Ave, Chicago, IL 60607.

## Abstract

Acetaminophen (APAP) overdose causes hepatic injury and is major contributor to acute liver injury cases. To investigate potential roles of gut microbiota in APAP-induced liver injury, C57BL/6 mice from Jackson (JAX) or Taconic (TAC) were challenged with APAP. TAC mice were more susceptible to APAP toxicity, and this disappeared upon co-housing of JAX and TAC mice. When the cecum contents from JAX and TAC mice were transplanted to germ-free mice, the mice that received TAC gut microbiota exhibited more significant hepatotoxicity after APAP administration. Non-targeted metabolomic analysis using portal vein serum and liver tissue of the mice led to identification of 19 metabolites the levels of which are associated with JAX or TAC gut microbiota. A gut bacteria-derived metabolite phenylpropionic acid (PPA) levels in cecum contents and blood were higher in mice harboring JAX gut microbiota. PPA supplementation in drinking water alleviated APAP-induced hepatotoxicity in TAC mice. This was accompanied by reduced hepatic protein levels of cytochrome P450 (CYP) 2E1, the enzyme responsible for APAP bioactivation to a toxic metabolite. This illustrates a gut microbe-liver interaction mediated by a gut bacteria-derived metabolite in modulating drug-induced liver injury.

## INTRODUCTION

Human gut carries ~100 trillions of bacteria that belong to >100 different species ^1^. Human gut microbes outnumber human cells by 3 times, and their gene pool size is 100-fold bigger than human genome ^1^. These gene products are involved in generation of thousands of chemically diverse small molecules from food ^2,3^, many of which are bioactive and mediate the intricate interactions between gut microbes and host. For example, short chain fatty acids (SCFA; up to ~100 mM in the gut ^4^) modulate host immune and metabolic systems ^5^ while aromatic amino acid metabolites regulate intestinal permeability and systemic immunity ^6,7^. Diet as well as metabolic gene contents among gut microbial communities govern the levels of gut microbial metabolites ^3,6,8,9^. Recent studies have demonstrated that the metabolic output of gut microbiota, given the same diet, can be controlled, for example by chemical inhibition of microbial enzymes mediating the metabolite production ^10,11^ or introduction of engineered bacteria strains into the complex microbial communities ^6,12^. Together, gut microbiota is an emerging and promising target that can be engineered to improve human health. What remains lacking is detailed understanding about the biological functions and the underlying molecular mechanisms for most gut bacteria-derived metabolites ^3,13^.

APAP is a commonly used analgesic but triggers fulminant acute liver failure upon overdose. APAP is the major cause of acute liver failure in the US ^14,15^. APAP is mainly eliminated as inactive sulfate or glucuronide conjugates into the urine, and a small fraction (<10%) of APAP is bioactivated to a toxic metabolite *N-*acetyl-*p*-benzoquinone imine (NAPQI) by cytochrome P450 (CYP) 2E1 (major), CYP3A4, and CYP1A2 ^16^. NAPQI is a strong oxidizing agent and interacts with intracellular antioxidant glutathione, leading to glutathione depletion in the liver. This subsequently leads to NAPQI binding to cellular components (e.g., proteins), oxidative stress, and cell necrosis.

Susceptibility to APAP-induced hepatotoxicity exhibits large interindividual variability ^17,18^. Multiple factors including age, food intake, genetic polymorphisms of APAP-metabolizing enzymes, and concurrent use of alcohol were shown to contribute to varying clinical outcomes after APAP overdose ^19–22^. Accumulating evidence suggests that altered gut microbiota may also impact the susceptibility to APAP hepatotoxicity. Metabolomics results of pre- and post-dose urinary metabolite profiles after the administration of a therapeutic dose of APAP suggest that gut bacteria metabolite *p*-cresol may competitively inhibit APAP sulfonation and potentially diverting APAP towards CYP-mediated bioactivation ^23^. Diurnal variations in gut microbiota abundance and composition were shown to be associated with altered susceptibility to APAP-induced hepatotoxicity ^24,25^. Also, a probiotic *Enterococcus lactis* IITRHR1 was shown to alleviate APAP hepatotoxicity potentially by its antioxidant effects ^26^.

In this study, we systematically investigated the roles of gut microbiota in APAP-induced hepatotoxicity and provided mechanistic understanding underlying host-gut microbiota interactions via a gut microbial metabolite, phenylpropionic acid (PPA).

## MATERIALS AND METHODS

### Co-housing of mice

Male or female C57BL/6 mice were purchased from Jackson laboratory (hereafter JAX; Sacramento, CA) or Taconic biosciences (hereafter TAC; Cambridge city, IN) and maintained in specified pathogen free state and fed a standard chow diet (Teklad 7012, Envigo). After one week of acclimation, mice were fasted overnight (14 h) and intraperitoneally injected with APAP (dissolved in PBS) or PBS. Food was returned to the mice after treatment. Mice were sacrificed 24 h after treatment. For cohousing, male C57BL/6 mice (4 weeks of age) from JAX or TAC were cohoused with mice from either the same or different vendors (4 mice/cage). After 4 weeks of cohousing, mice were fasted overnight and sacrificed. All animal procedures were approved by the Institutional Animal Care and Use Committee in the University of Illinois at Chicago.

### Gut microbiota transplantation

Cecum materials were collected from JAX or TAC mice (8-9 weeks of age; housed at University of Illinois at Chicago), pooled, and suspended in PBS (100 mg/mL). The mixtures were homogenized and centrifuged to remove debris. The supernatants (200 μL) were orally administered to male C57BL/6 germ-free (GF) mice (8-12 weeks of age; University of Chicago). Mice were fed a standard chow diet (Lab diet^®^ 6F 5K67, Lab diet). After 4 weeks, mice were fasted overnight and sacrificed. All animal procedures involving GF mice were approved by the University of Chicago Animal Care and Use Committee.

### PPA supplementation

PPA were dissolved in drinking water at the final concentration of 0.4% (w/v), followed by pH adjustment. C57BL/6 mice from TAC (6-7 weeks of age) had free access to drinking water. After 4 weeks, mice were fasted overnight and sacrificed.

### Liver histology and measurement of ALT

Liver tissues were fixed with 10% neutral buffered formalin. Fixed liver tissues were embedded in paraffin. Sections were cut and stained with hematoxylin and eosin (H/E). Serum ALT levels were measured using ALT kit (A7526-150, Pointe Scientific, MI) according to the manufacturer’s instruction.

### Measurement of hepatic glutathione

Hepatic glutathione levels were measured in the liver homogenate using a commercially available kit from Dojindo molecular technologies (Rockville, MD). Briefly, liver tissues were homogenized in 5% sulfosalicylic acid solution. The homogenates were centrifuged at 8,000*g* for 10 min (4°C). The supernatant was stored at −80°C until assay. Assay was performed using manufacturer’s instructions.

### Fecal microbiota genomic DNA extraction

Fecal pellets were collected before fasting, snap-frozen, and stored at −80°C until analysis. Genomic DNA was extracted using a Tissue DNA Purification Kit, implemented on a Maxwell® 16 automated extraction system (Promega). Genomic DNA quantity was assessed using a Qubit 2.0 fluorometer with the dsDNA BR Assay (Life Technologies, Grand Island, NY).

### 16S rRNA gene amplicon sequence libraries and data analysis

The V4 region of the 16S rRNA gene was amplified and sequenced at the High-Throughput Genome Analysis Core at Argonne National Laboratory. Briefly, PCR amplicon libraries targeting the 16S rRNA encoding gene present in metagenomic DNA are produced using a barcoded primer set adapted for the Illumina HiSeq2000 and MiSeq ^27^. DNA sequence data is then generated using Illumina paired-end sequencing at the Environmental Sample Preparation and Sequencing Facility (ESPSF) at Argonne National Laboratory. Specifically, the V4 region of the 16S rRNA gene (515F-806R) is PCR amplified with region-specific primers that include sequencer adapter sequences used in the Illumina flowcell ^27,28^. The forward amplification primer also contains a twelve-base barcode sequence that supports pooling of up to 2,167 different samples in each lane ^27,28^. Each 25 μL PCR reaction contains 9.5 μL of MO BIO PCR Water (Certified DNA-Free), 12.5 μL of QuantaBio’s AccuStart II PCR ToughMix (2x concentration, 1x final), 1 μL Golay barcode tagged Forward Primer (5 μM concentration, 200 pM final), 1 μL Reverse Primer (5 μM concentration, 200 pM final), and 1 μL of template DNA. The conditions for PCR are as follows: 94°C for 3 minutes to denature the DNA, with 35 cycles at 94°C for 45 s, 50°C for 60 s, and 72°C for 90 s; with a final extension of 10 min at 72°C to ensure complete amplification. Amplicons are then quantified using PicoGreen (Invitrogen) and a plate reader (Infinite ® 200 PRO, Tecan). Once quantified, volumes of each of the products are pooled into a single tube so that each amplicon is represented in equimolar amounts. This pool is then cleaned up using AMPure XP Beads (Beckman Coulter), and then quantified using a fluorometer (Qubit, Invitrogen). After quantification, the molarity of the pool is determined and diluted down to 2 nM, denatured, and then diluted to a final concentration of 6.75 pM with a 10% PhiX spike for sequencing on the Illumina MiSeq. Amplicons are sequenced on a 151bp × 12bp × 151bp MiSeq run using customized sequencing primers and procedures ^27^.

Quantitative Insights into Microbial Ecology 2 (QIIME2) software (https://qiime2.org/) was used to analyze the sequencing data. Forward, reverse and barcodes sequences were imported into an artifact type of EMPPairedEndSequences and demultiplexed ^29^. Demultiplexed reads were denoised with DADA2 ^30^ generating operational taxonomic units (OTU) table. Taxonomic annotations for each OTU were generated with the taxonomy classifier trained on the Greengenes18_3 97% OTUs where the sequences have been trimmed to include bases from the region of the 16S that was sequenced in this analysis (the V4 region, bound by the 515F/806R primer pair) ^31^. Taxonomic and OTU abundance data were generated for all phyla, classes, orders, families, genera, and species. Nonmetric multidimensional scaling (NMDS) plot, analysis of similarities (ANOSIM) and Shannon-index was created or calculated using the R package ‘ggplot’ and ‘vegan’. False discovery rates (FDRs) of the multiple comparisons were estimated for each taxon based on the *p*-values resulting from Spearman correlation estimates using R package ‘multtest’.

### Measurement of APAP-sulfate and *p*-cresol sulfate in the liver

Liver tissues (~50 mg) were homogenized in 450 μL of ice-cold methanol followed by centrifugation at 16,000*g* for 10 min at 4°C. Supernatants were filtered through 0.22 μm pore Ultrafree-MC filter and stored at −80°C until further use. Filtrates (50 μL) were mixed with 200 μL of methanol containing internal standards (mixtures of APAP-sulfate-D_3_ and *p*-cresol-sulfate-D_7_) and centrifuged at 16,000*g* for 10 min at 4°C. The supernatants were used for liquid chromatography tandem mass spectrometer (LC-MS/MS) analysis.

The concentrations of APAP-sulfate and *p*-cresol sulfate were measured in positive and negative ion modes, respectively, using LC-MS/MS. The separation was performed on an Atlantis T3 column (3 μm, 100 mm × 3 mm, i.d., Waters Corp., Milford, MA). Chromatographic separation was performed using 0.1 % formic acid in water (solvent A) and 0.1 % formic acid in acetonitrile (solvent B) at a flow rate of 0.4 mL/min. The gradient elution profile used is as follows: 95% A for 3 min, 95% to 10% A over 3 min, 10% A for 3 min, 10% to 95% A over 0.5 min, and 95% A for 5 min. The detection and quantification of analytes were accomplished by multiple reactions monitoring of 232.1/152.1 for APAP-sulfate; 235.1/155.1 for APAP-sulfate-D_3_; 328.1/152.1 and 187/107 for *p*-cresol sulfate; and 194/114 for *p*-cresol sulfate-D_7_.

### Measurement of short chain fatty acids (SCFAs) in mouse cecum contents

Mouse cecum content (~50 mg) was mixed with 250 μL of 50% aqueous acetonitrile, vortexed for 10 s and centrifuged at 16,000*g* for 15 min at 4°C. Supernatant (200 μL) was collected and analyzed. Mixed standard solutions containing equal amount of individual SCFA at desired concentration were made in 50% acetonitrile. For derivatization, 40 μL of each standard solution or of each the supernatant were mixed with 20 μL of 10 mM 3-NPH in 50% acetonitrile and 20 μL of 120 mM EDC in the same solution, incubated for 30min at 40°C, and 10% acetonitrile (1 mL) was added. Diluted reaction solution was mixed with pre-made stable isotope labeled standards. The LC-MS/MS analysis was performed on AB Sciex Qtrap 5500 coupled to Agilent UPLC/HPLC system. All samples are analyzed by Agilent poroshell 120 EC-C18 Column (100Å, 2.7 μm, 2.1 mm X 100 mm). Chromatographic separation was performed using a gradient of solvent A (water, 0.1% formic acid) and solvent B (acetonitrile, 0.1% formic acid) was applied as: 0 min, 30% solvent B; increase solvent B to 100% over 4 min; maintain solvent B at 100% for 5 min at a flow rate of 400 μL/min. The column was equilibrated for 3 min at 30% solvent B between the injections. The column temperature was 25°C and the autosampler was kept at 4°C. The detection and quantification of analytes were accomplished by multiple reactions monitoring of *m/z* 136.1/94.1 for acetate; 142.1/100.1 for acetate-^13^C_6_; 150.1/94.1 for propionate; 156.1/100.1 for propionate-^13^C_6_; 164.1/94.1 for butyrate; and 170.1/100.1 for butyrate-^13^C_6_.

### Non-targeted metabolomics

Portal vein serum and liver tissues were collected before APAP administration from conventional JAX and TAC mice as well as JAX_GF_ and TAC_GF_ mice (n=5 mice/group). Respective samples from GF mice (n=3/group) were also included as a control. Samples were immediately snap frozen in liquid nitrogen and stored at −80°C until analysis. Samples for metabolomic analysis were prepared using the automated MicroLab STAR® system from Hamilton Company and analyzed via UPLC-MS/MS by Metabolon, Inc. Proteins were precipitated with methanol under vigorous shaking for 2 min (Glen Mills GenoGrinder 2000) followed by centrifugation. The resulting extracts were divided into five fractions: two for analysis by two separate reverse phase (RP)/UPLC-MS/MS methods with positive ion mode electrospray ionization (ESI), one for analysis by RP/UPLC-MS/MS with negative ion mode ESI, one for analysis by HILIC/UPLC-MS/MS with negative ion mode ESI, and one sample was reserved for backup. Samples were placed briefly on a TurboVap® (Zymark) to remove the organic solvent. The sample extracts were stored overnight under nitrogen before preparation for analysis. For UPLC-MS/MS analysis, all methods utilized a Waters ACQUITY UPLC and a Thermo Scientific Q-Exactive high resolution/accurate mass spectrometer interfaced with a heated electrospray ionization (HESI-II) source and Orbitrap mass analyzer operated at 35,000 mass resolution. The sample extracts were dried and reconstituted in solvents compatible to each of the four methods. Each reconstitution solvent contained a series of standards at fixed concentrations to ensure injection and chromatographic consistency. One aliquot was analyzed using acidic positive ion conditions, chromatographically optimized for more hydrophilic compounds. In this method, the extract was gradient eluted from a C18 column (Waters UPLC BEH C18-2.1×100 mm, 1.7 μm) using water and methanol, containing 0.05% perfluoropentanoic acid (PFPA) and 0.1% formic acid. Another aliquot was also analyzed using acidic positive ion conditions which was chromatographically optimized for more hydrophobic compounds. In this method, the extract was gradient eluted from the same afore mentioned C18 column using methanol, acetonitrile, water, 0.05% PFPA and 0.01% formic acid and was operated at an overall higher organic content. Another aliquot was analyzed using basic negative ion optimized conditions using a separate dedicated C18 column. The basic extracts were gradient eluted from the column using methanol and water containing 6.5 mM ammonium bicarbonate at pH 8. The fourth aliquot was analyzed via negative ionization following elution from a HILIC column (Waters UPLC BEH Amide 2.1×150 mm, 1.7 μm) using a gradient consisting of water and acetonitrile with 10 mM ammonium formate, pH 10.8. The MS analysis alternated between MS and data-dependent MS^n^ scans using dynamic exclusion. The scan range covered 70-1000 *m/z*. Peaks were quantified using area-under-the curve. Detected metabolites were identified by comparison to library entries of purified standards. Following normalization to serum sample volume or liver tissue weight, log transformation and imputation of missing values, Welch’s two-sample t-test was used to identify metabolites that significantly different between experimental groups.

### Measurement of 3-phenylpropionic acid in mouse cecum and serum

Mouse cecum contents (~50 mg) were mixed with 250 μL of 50% aqueous acetonitrile, vortexed for 10 s and centrifuged at 16,000*g* for 15 min at 4°C. Supernatants were collected, diluted with 50% acetonitrile by 20 fold, and mixed with ice-cold acetonitrile containing *trans*-cinnamic acid (tCA)-d_6_ (100 ng/mL) as an internal standard followed by centrifugation at 16,000*g* for 10 min at 4°C. Supernatants were filtered through 0.22 μm pore Ultrafree-MC filter and used for LC-MS/MS analysis. PPA-d_9_ was used as a surrogate standard to construct the calibration curve for the quantification of PPA ^32^.

Mouse serum collected from portal vein or heart was diluted with double distilled water (DDW) by 3 fold and mixed with ice-cold acetonitrile containing 100 ng/mL of tCA-d_6_ followed by centrifugation at 16,000*g* for 10 min at 4°C. Supernatants were dried and reconstituted with 70:30 mixture of water (0.01% acetic acid) and acetonitrile (0.01% acetic acid).

The levels of PPA in the cecum filtrates or serum were measured using LC-MS/MS in negative ion mode. The separation was performed on Atlantis T3 column (3 μm, 100 mm × 3 mm, i.d., Waters Corp., Milford, MA). Chromatographic separation was performed using 0.01% acetic acid in water (solvent A) and 0.01% acetic acid in acetonitrile (solvent B) at a flow rate of 0.4 mL/min. The gradient elution profile used is as follows: 70% solvent A for 0.5 min, 70% to 10% solvent A over 5 min, 10% solvent A for 4 min, 10% to 70% solvent A over 0.1 min, and 70% solvent A for 5.4 min (15 min analysis in total). The detection and quantification of analytes were accomplished by multiple reactions monitoring of *m/z* 149.0/105.0 for PPA; 158.0/114.2 for PPA-d_9_; 147.0/103.2 for tCA; and 152.9/109.1 for tCA-d_6_. tCA was not detected in either cecum filtrates or serum.

### Isolation of mouse liver microsome

Liver tissues (~100 mg) were homogenized in 1000 μL of ice-cold buffer 1 [225 mM mannitol, 75 mM sucrose, 30 mM Tris, 0.5 m EDTA (pH 7.4), protease inhibitor (Roche 04 693 123 004), phosphatase inhibitor (Roche 04 906 837 001)] followed by centrifugation at 800*g* for 5 min at 4°C. Supernatants were collected and centrifuged at 800*g* for 5 min at 4°C for 4 more times to remove unbroken cells and nuclei. Supernatants were collected and centrifuged at 9,000*g* for 10 min at 4°C. Supernatants were collected and centrifuged at 20,000*g* for 30 min at 4°C to remove peroxisome pellets. Supernatants were collected and centrifuged at 100,000*g* for 60 min at 4°C. Supernatants were discarded and microsomal pellets were resuspended with buffer and centrifuged at 100,000*g* for 30 min at 4°C. Pellets were resuspended with microsome buffer (170 mM KCl, 50 mM KPO_4_, pH 7.4 containing protease inhibitor) and stored at −80°C until use.

### Isolation of mouse liver mitochondria

Liver mitochondria was isolated using differential centrifugation followed by Percoll density gradient ^33^. Liver tissue (~850 mg) was homogenized in 6 ml of buffer 1 [225 mM Mannitol, 75 mM sucrose, 30 mM Tris (pH 7.4), 0.5 mM EDTA] containing protease and phosphatase inhibitors (Roche 04 693 123 004 and 04 906 837 001). Supernatant was centrifuged at 740*g* for 5 min at 4°C. Supernatant was collected and repeated centrifugation and collection for additional 3 times. Supernatant was centrifuged at 9,000*g* for 10 min at 4°C. Supernatant was collected for cytosol and microsome fractionation. Crude mitochondria pellet was resuspended with buffer 2 [225 mM mannitol, 75 mM sucrose, 30 mM Tris buffer (pH 7.4), 0.5% BSA] and centrifuged at 10,000*g* for 10 min at 4°C. Pellets were resuspended with buffer 3 [225 mM mannitol, 75 mM sucrose and 30 mM Tris buffer (pH 7.4)] and centrifuged at 10,000*g* for 10 min at 4°C. Pellets were resuspended with mitochondria resuspension buffer [MRB, 250 mM mannitol, 5 mM HEPES (pH 7.4) and 0.5 mM EDTA], added on top of 30% (v/v) Percoll solution [30% Percoll : 70% buffer consisting of 225 mM mannitol, 25 mM HEPES (pH 7.4), 1mM EDTA] and centrifuged at 95,000*g* for 30 min at 4°C. Pure mitochondria band was localized at the bottom of tube and collected. Pure mitochondria solution was diluted with MRB by 10 folds and centrifuged for 10 min at 4°C. Pure mitochondria pellet was resuspended with MRB with protease and phosphatase inhibitor.

For concomitant isolation of cytosol and microsome, the supernatant containing cytosol and microsome fraction was centrifuged for 30 min at 4°C. Supernatant was collected and centrifuged at 100,000*g* for 1 h at 4°C and supernatant was collected as the cytosol fraction. Microsomal pellet was resuspended with microsome buffer [0.17 M KCl, 50 mM KPO_4_ (pH 7.4)] and centrifuged again at 100,000*g* for 30 min at 4°C. Pellet was resuspended with the buffer containing protease and phosphatase inhibitor.

### Measurement of APAP bioactivation

The extent of APAP bioactivation in hepatic microsome was measured as previously described with modifications ^34,35^. Briefly, APAP (at a final concentration of 1 mM) was incubated with microsome (2.5 μg) in mixtures (20 μL total volume) containing GSH (10 mM) and a NADPH-generating system [0.5 mM NADP^+^, 1 mM MgCl_2_, 0.2 U/L isocitrate dehydrogenase, and 0.5 mM isocitric acid in 100 mM Tris buffer (pH 7.4)]. The reaction was stopped after 30 min by adding 100 μL of ice-cold acetonitrile containing 100 ng/ml APAP-d_4_ (internal standard). The mixtures were centrifuged at 12,000*g* for 20 min (4°C), and APAP-GSH level in the supernatant was measured using LC-MS/MS in the positive ionization mode (QTrap 5500; Applied Biosystems, ON, Canada). The separation was performed on an Atlantis T3 column (3 μm, 100 mm × 3 mm, i.d., Waters Corp., Milford, MA). Chromatographic separation was performed using 0.1 % formic acid in water (solvent A) and 0.1 % formic acid in acetonitrile (solvent B) at a flow rate of 0.4 mL/min. The gradient elution profile used is as follows: 95% A for 3 min, 95% to 10% A over 3 min, 10% A for 3 min, 10% to 95% A over 0.5 min, and 95% A for 5 min. The detection and quantification of analytes were accomplished by multiple reactions monitoring of *m/z* 457.1/140.0 for APAP-GSH and 156.1/114.0 for APAP-d_4_.

### Measurement of CYP2E1 activity

Chlorzoxazone at a final concentration of 100 μM was incubated with mouse liver microsome (25 μg) in mixtures (100 μL total volume) containing the NADPH-generating system, along with 1 mM of PPA or *p*-nitrophenol. The reaction was stopped by adding 200 μL of acetonitrile containing 0.4 mM 2-benzoxazolinone (internal standard) after 30 min. Precipitated proteins were removed by centrifugation (12,000*g*, 4°C, 20 min), and the supernatant was analyzed using liquid chromatography tandem mass spectrometer (LC-MS/MS) in the negative ionization mode (QTrap 5500; Applied Biosystems, ON, Canada). The separation was performed on a XTerra MS C18 column (2.5 μm, 50 mm × 2.1 mm, i.d., Waters Corp., Milford, MA). The HPLC system (Agilent 1200, Santa Clara, CA) was operated isocratically at a flow rate of 0.3 mL/min. A mixture of acetonitrile (0.1% formic acid) (solvent A) and 0.1 mM ammonium formate (0.1% formic acid) (solvent B) was used as the mobile phase (A:B, 20:80). The detection and quantification of analytes were accomplished by multiple reactions monitoring with the transitions of *m/z* 183.9/119.9 for 6-hydroxychlorzoxazone; 167.9/132.1 for chlorzoxazone; and 133.9/64.9 for 2-benzoxazolinone.

### Measurement of APAP-protein adduct in the liver

APAP-protein adduct was measured as previously described with modifications^36,37^. Briefly, liver tissue (~50 mg) was homogenized in 10 mM sodium acetate buffer (500 μL, pH 6.5) and filtered through Bio-spin 6 column (Bio-Rad, Hercules, CA) prewashed with 10 mM sodium acetate buffer (pH 6.5) to remove free APAP and APAP metabolites. The filtrate was incubated with protease type XIV enzyme in water (8 U/mL; 50 μL) at 37 °C for 24 h. Ice-cold acetonitrile (800 μL) containing APAP-d_4_ as an internal standard (20 ng/ml) were added and centrifuged at 16,000*g* for 10 min at 4 °C. Supernatant (800 μL) was dried and reconstituted in mixture of 0.1% formic acid water and 0.1% formic acid acetonitrile (95:5) followed by centrifugation at 16,000*g* for 10 min at 4 °C. The supernatants were filtered through Ultrafree-MC centrifugal filter to remove large particle, and APAP-cysteine (APAP-CYS) in the filtrate was measured using LC-MS/MS in positive ion mode. The separation was performed on an Atlantis T3 column (3 μm, 100 mm × 3 mm, i.d., Waters Corp., Milford, MA). Chromatographic separation was performed using 0.1 % formic acid in water (solvent A) and 0.1 % formic acid in acetonitrile (solvent B) at a flow rate of 0.4 mL/min. The gradient elution profile used is as follows: 95% A for 3 min, 95% to 10% A over 3 min, 10% A for 3 min, 10% to 95% A over 0.5 min, and 95% A for 5 min. The detection and quantification of analytes were accomplished by multiple reactions monitoring of *m/z* 270.9/139.8 for APAP-CYS and 155.9/114.1 for APAP-d_4_. Protein level of liver homogenate filtrate was determined using a standard bicinchoninic acid assay and APAP-CYS level was normalized to total protein in the liver homogenates.

### Western blot

Hepatic microsomes were separated using sodium dodecyl sulfate polyacrylamide gel electrophoresis (SDS-PAGE) and transferred to polyvinylidene difluoride membrane. Membrane was blocked using 5% skim milk in Tris-buffered saline, 0.1% Tween 20 at room temperature for one hour and incubated with antibodies against calnexin (Abcam, ab22595), CYP2E1 (Abcam, ab28146), CYP1A2 (Chemicon, Mab10035), voltage-dependent anion channel (VDAC; Thermo, PA1-954A), or α-tubulin (Calbiochem, CP06) overnight at 4 °C. On the next day, the membrane was incubated with secondary antibody (1:10000) for one hour, and proteins were detected using chemiluminescence (Pierce Biotechnology). The intensity of protein bands was quantified using Image J.

### RNA sequencing

Total RNA was isolated from liver tissues (n=4-5 mice/group) using Trizol (ThermoFisher Scientific). Concentration was measured by nanodrop (ThermoFisher Scientific) at 260 nm. RNA samples (n=3/group) from 4 groups of mice were used for RNA sequencing. From 5 μg of total RNA, mRNA was isolated using NEBNext Poly(A) mRNA Magnetic Isolation Module (cat. No. E7490, New England Biolabs, Ipswich, MA) following manufacturer’s instructions. Fragmentation of RNA followed by reverse transcription and second strand cDNA synthesis was performed using NEBNext Ultra RNA Library Prep Kit for Illumina (cat. No. E7530, NEB). The resulting double-stranded cDNA’s were further processed to DNA sequencing libraries using ThruPLEX DNA-seq 12S Kit (Rubicon Genomics, Ann Arbor, MI). Libraries were selected by gel purification for an average size of 350 bp. Each purified library was quantified using a Qubit fluorometer (Life Technologies) and equal amounts of each library were pooled and sequenced on the Illumina HiSeq 2500. All RNA sequencing reads were aligned to mm9 using Tophat. The mRNA abundance was expressed as Counts Per Million (cpm).

### Quantitative real-time (qRT)-PCR

Total RNA was isolated from liver tissues using Trizol (Life Technologies) and used as template for the synthesis of complementary DNA using High Capacity cDNA Archive Kit (Applied Biosystems, Foster City, CA). Using the cDNA as template, qRT-PCR was performed using the following primers from Integrated DNA Technologies (California, USA): Cyp2e1 (Mm.PT.58.9617541), Gclc (Mm.PT.58.30656560), and Gapdh (Mm.PT.39a.1).

### Statistical analysis

All data were expressed as mean ± standard deviation (SD). Comparisons between groups were made by using the Student’s *t*-test or one-way ANOVA via GraphPad Prism 7. *P*-value less than 0.05 was considered significant. Differential expression statistics (fold-change and *p*-value) for gut microbiome and transcriptome were computed using the exactTest function in edgeR ^38,39^ on raw expression counts obtained from quantification. *P*-values were adjusted for multiple testing using the false discovery rate (FDR) correction of Benjamini and Hochberg ^40^. FDR value less than 0.05 was considered significant.

## RESULTS

### Differential gut microbiota modulates susceptibility to APAP toxicity

Mice raised in different environments are known to harbor distinct gut microbiota composition^41,42^. To determine whether differential gut microbiota modulates susceptibility to APAP toxicity, male C57BL/6 mice from two different vendors [Jackson (JAX) and Taconic (TAC)] were challenged with APAP. Considering the inter-laboratory variations in the susceptibility threshold to fulminant APAP-induced hepatotoxicity, APAP dose that exhibits prominent hepatotoxicity was first determined. Mice were treated with 100, 200 or 300 mg/kg APAP (or PBS) via intraperitoneal injection and sacrificed 24 h after treatment. Serum levels of ALT were measured as a marker of liver damage. All mice in the PBS or 100 mg/kg groups and about half of the mice in the 200 mg/kg group did not exhibit hepatotoxicity (Fig S1A). Accordingly, the APAP dose of 300 mg/kg was chosen for the study. At this dose, the extent of hepatotoxicity (i.e., serum ALT level) was much greater in TAC than JAX mice (Fig S1B). A similar trend was observed in female mice although the severity of hepatotoxicity was less in female mice (Fig S1B), consistent with the previous reports^43^. Subsequent studies for APAP-induced hepatotoxicity were performed using male mice.

To examine whether differential gut microbiota contributes to the differences between JAX and TAC mice in APAP hepatotoxicity, the mice were cohoused for 4 weeks (a process known to assimilate gut microbiota of mice) and challenged with APAP. This created two additional groups of mice, namely coJAX (i.e., JAX mice cohoused with TAC) and coTAC (i.e., TAC mice cohoused with JAX). At the end of 4-week cohousing, fecal samples were collected from each mouse, and gut microbiome was analyzed by 16S rRNA gene amplicon sequencing. Nonmetric multidimensional scaling (NMDS) plot showed that JAX mouse group was separated from TAC mice (*p*=0.001, ANOSIM) and while coJAX and coTAC groups clustered together (*p* = 0.149, ANOSIM) (Fig 1A), suggesting that JAX and TAC mice indeed had distinct gut microbiota composition and co-housing abrogated the difference. Similarly, the difference in Shannon-index (an indicator of species richness or α-diversity) between JAX and TAC mice disappeared upon cohousing (Fig S1C). Importantly, the difference in the serum ALT levels and hepatic necrosis between JAX and TAC mice disappeared upon co-housing (Fig 1B and Fig S1E).

**Fig 1.**
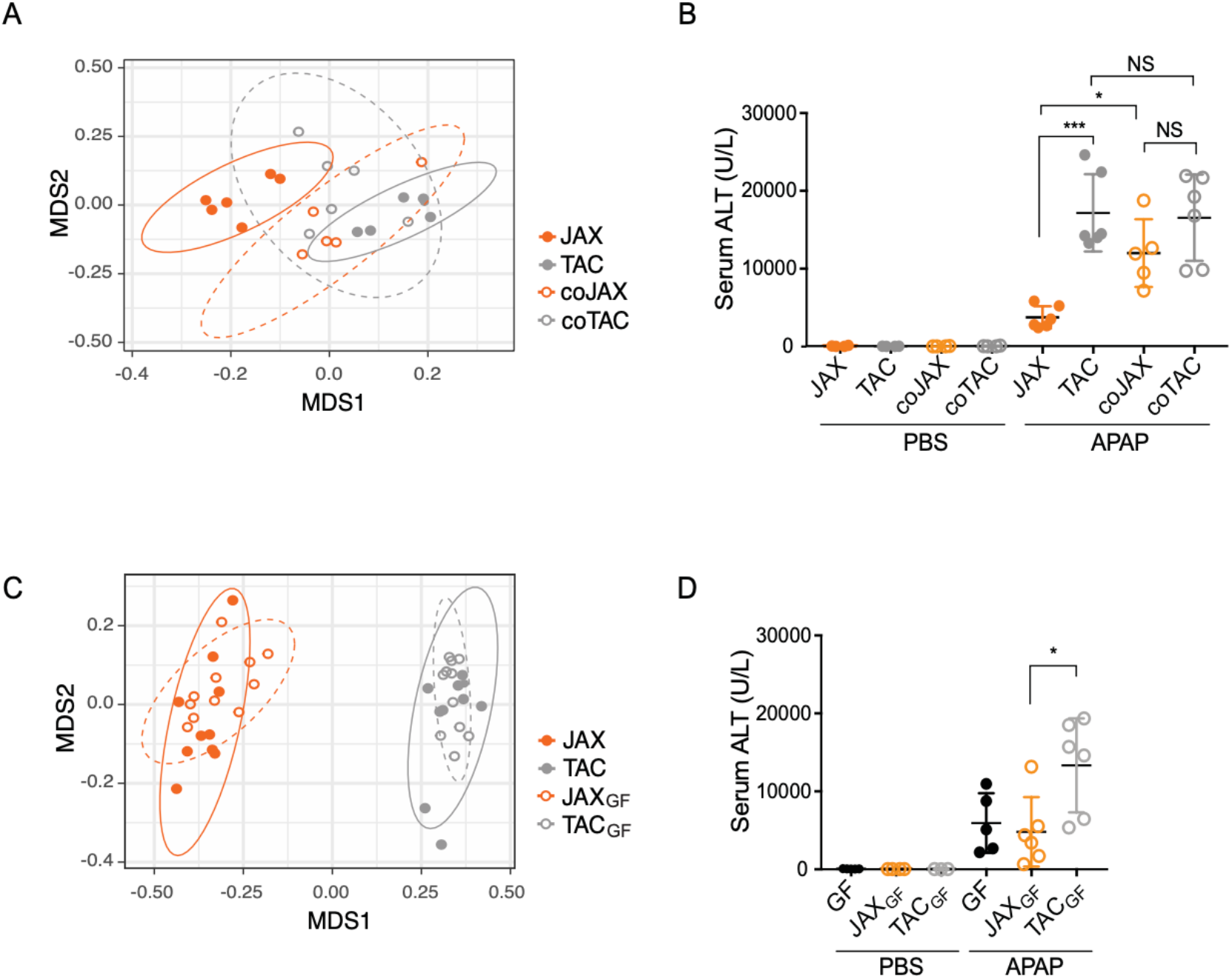
Differential gut microbiota modulates susceptibility to APAP-induced hepatotoxicity. **A and B**. C57BL/6 mice from JAX or TAC companies were co-housed with mice from the same or the other vendor for 4 weeks. (A) Gut microbiome was analyzed by 16S rRNA gene amplicon sequencing using fecal samples collected at the end of co-housing. Nonmetric multidimensional scaling (NMDS) plot is shown. (B) APAP (300 mg/kg or PBS, intraperitoneal injection) was administered after overnight fasting. Mice were sacrificed after 24 h, and serum ALT levels were measured as a measure of hepatotoxicity. **C and D**. Pooled cecum contents from JAX or TAC mice were orally administered to GF C57BL/6 mice. After 4 weeks, fecal samples were collected, and the mice were challenged with APAP (300 mg/kg or PBS, intraperitoneal injection). (C) NMDS plot based on 16S rRNA gene amplicon sequencing of the fecal samples is shown. APAP (300 mg/kg or PBS, intraperitoneal injection) was administered after overnight fasting. (D) Serum ALT levels at 24 h post-dosing are shown. All data are shown as mean ± S.D.

Genetic drifts between C57BL/6 mice from JAX and TAC have been reported^44^. To examine the roles of differential gut microbiota in APAP toxicity in the mice of the same genetic background, the gut microbiota from JAX or TAC mice was transplanted to GF C57BL/6 mice, followed by APAP challenge. To this end, pooled cecum contents from JAX or TAC mice were orally administered to GF mice, followed by 4-week conventionalization (generating JAX_GF_ and TAC_GF_ groups, respectively). Fecal microbiome analysis revealed that gut microbiota of JAX_GF_ and TAC_GF_ mice were distinct (*p*=0.001, ANOSIM) and co-clustered with their respective donor groups (Fig 1C). Also, Shannon-index of JAX_GF_ and TAC_GF_ were similar to those of donor mice, indicating that cecum material transplantation successfully established the donor’s microbiome in GF mice (Fig S1D). When challenged with APAP, TAC_GF_ mice had higher serum ALT levels and greater hepatic necrosis compared to JAX_GF_, recapitulating the phenotype observed in conventional JAX and TAC mice (Fig 1D and Fig S1F), indicating that differential gut microbiota modulates APAP-induced hepatotoxicity.

### Abundances of certain gut microbial taxa are associated with APAP toxicity

To understand how differential gut microbiota alters APAP-induced liver injury, the bacteria compositions in feces collected before APAP dosing were compared between JAX and TAC gut microbiota. At the phylum level, *Bacteroidetes* and *Firmicutes* accounted for the majority of bacteria (80-90%), consistent with a previous report for mouse gut microbiota ^45^. JAX and JAX_GF_ mice exhibited significantly higher abundances of *Verrucomicrobia*, and TAC and TAC_GF_ had higher *Deferribacteres* (Fig S2A). At the family level, JAX and JAX_GF_ gut microbiota was associated with higher abundances of *Verrucomicrobiaceae* while TAC and TAC_GF_ gut microbiota with higher abundances of *Porphyromonadaceae*, *Deferribacteraceae, Peptostreptococcaceae* and *Enterobacteriaceae* (Fig S2B), in part consistent with a previous report ^46,47^. At the species level, unclassified *Parabacteroides*, *Lactobacillus salivarius, Mucispirillum schaedleri,* and *Butyricicoccus pullicaecorum* were more abundant in TAC and TAC_GF_ gut microbiota, and *Akkermansia muciniphila* was less abundant (Table S1).

We further examined whether the relative abundances of any bacteria taxa were correlated with serum ALT level (at 24 h post-dosing). Fecal abundances of unclassified *Parabacteroides* and *Lactobacillus salivarius* were positively associated with serum ALT level (Fig 2A) in the mice from co-housing experiment. The abundance of unclassified *S24-7* was negatively associated with APAP toxicity. When these correlations were tested in gut microbiota-transplanted mice, similar trends were observed (Fig 2B) although they did not reach statistical significance potentially due to a relatively small sample size. Of note, fecal *A. muciniphila* or *B. pullicaecorum* abundances were found not correlated with serum ALT levels because its relative abundances in co-housed mice were similar to those in JAX mice, yet these mice exhibited greater APAP toxicity than JAX mice (Fig S2C). Additionally, fecal *M. schaedleri* abundances were not correlated with ALT levels due to large inter-mouse variability in the relative bacterial abundance.

**Fig 2.**
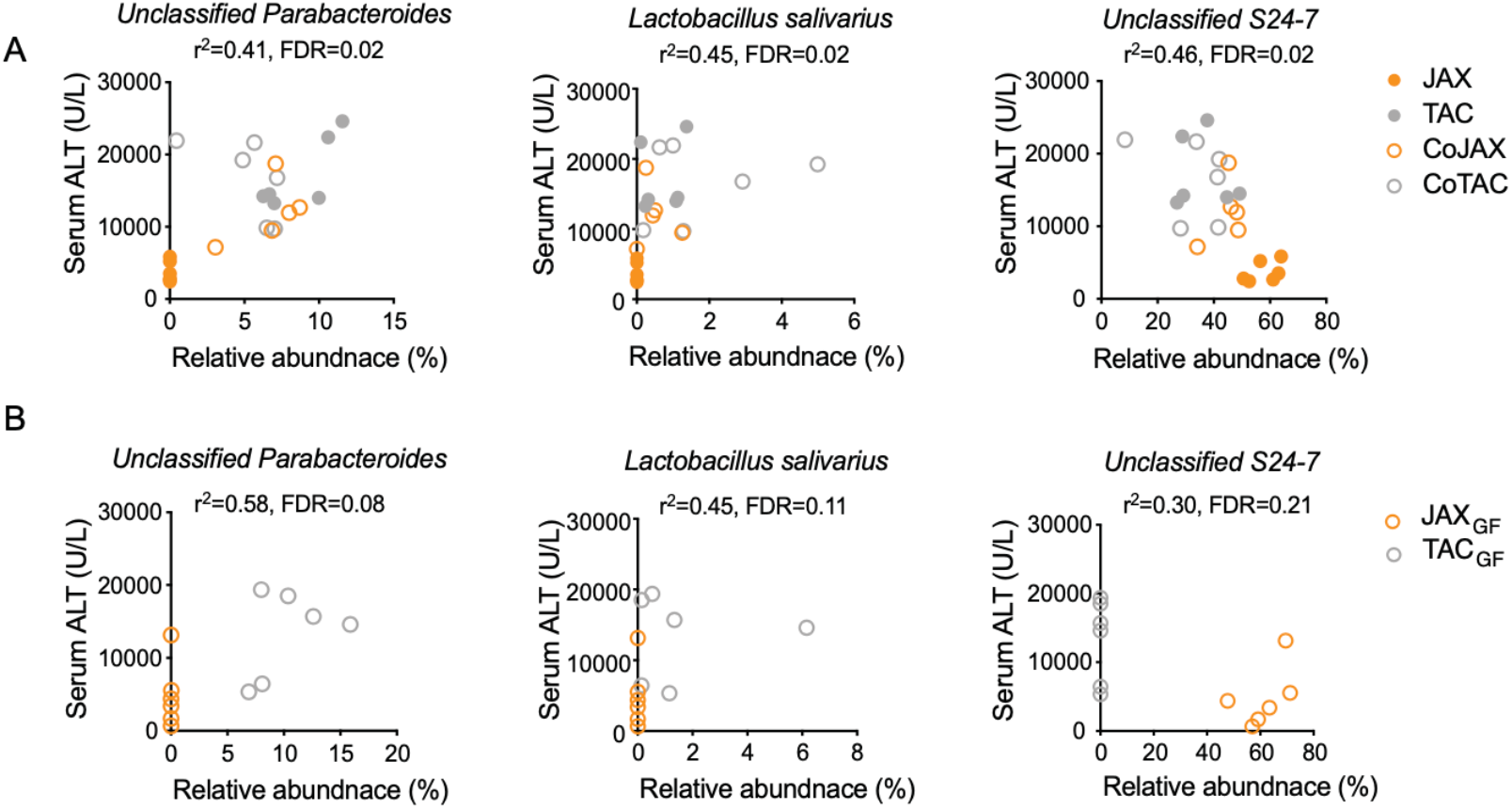
Fecal abundances of certain bacteria genus are associated with susceptibility to APAP-induced hepatotoxicity. **A.** C57BL/6 mice from JAX or TAC companies were co-housed with mice from the same or the other vendor for 4 weeks. Gut microbiome was analyzed by 16S rRNA gene amplicon sequencing using fecal samples collected at the end of co-housing. **B**. Pooled cecum contents from JAX or TAC mice were orally administered to GF C57BL/6 mice. After 4 weeks, fecal samples were collected and subjected to 16S rRNA gene amplicon sequencing.

### Gut microbiota-derived metabolite PPA reduces APAP toxicity

Gut microbiota produces thousands of chemically diverse small molecules, many of which are bioactive and influence host health and diseases^2,3^. Gut bacteria metabolite *p*-cresol may competitively inhibit APAP sulfonation, potentially diverting APAP towards CYP-mediated bioactivation ^23^. Basal hepatic *p*-cresol sulfate levels were higher in TAC than JAX mice (Fig S3A). On the other hand, hepatic APAP-sulfate levels (measured at 15 min after APAP dosing) were similar between TAC and JAX, and about 1000-fold higher than the *p*-cresol sulfate level (Fig S3B), suggesting that at a toxic dose of APAP, the competition of APAP sulfonation by *p*-cresol is unlikely.

Short chain fatty acids (SCFA) are major products of bacterial fermentation and modulate multiple aspects of host physiology ^48–50^. The cecum levels of major SCFAs, acetate and butyrate, were similar between JAX and TAC as well as JAX_GF_ and TAC_GF_ mice (Fig S2C). Propionate levels differ between JAX and TAC, but not in gut microbiota-transplanted mice. These results suggest minor roles of these SCFAs, if any, in modulating APAP-induced hepatotoxicity.

For *de novo* identification of gut microbiota-associated metabolites associated with altered susceptibility to APAP hepatotoxicity, portal vein serum and liver tissues collected from JAX and TAC mice as well as JAX_GF_ and TAC_GF_ mice were subjected to non-targeted metabolomic profiling using LC-MS/MS. Portal vein serum and liver tissue samples from GF mice were included as controls. A total of 789 and 669 compounds of known identity were detected in the liver and serum samples, respectively. A majority (i.e., 568 compounds) were detected in both liver and serum samples. Principal component analysis (PCA) of liver samples revealed good separation along component 1 between conventional (JAX and TAC) mouse and gut microbiota-transplanted (JAX_GF_ and TAC_GF_) mouse groups (Fig 3A), likely reflecting the effects of different facilities (i.e., University of Illinois at Chicago and University of Chicago for conventional mice and gut microbiota-transplanted mice, respectively) on metabolomes. Additional segregation was observed for JAX and TAC groups, as well as for GF, JAX_GF_, and TAC_GF_ groups. Hierarchical clustering analysis revealed separation of GF group (as expected), with further separation dictated by colonization status (i.e. conventionally raised vs. microbiota-transplanted mice. JAX_GF_ and TAC_GF_ samples were intermixed but showed group-related sub-clustering. Similar results were obtained from portal vein serum samples (Fig 3B). Taken together, these analyses indicate that in addition to GF status, the differences in microbiota structure dictate host metabolomic profiles.

**Fig 3.**
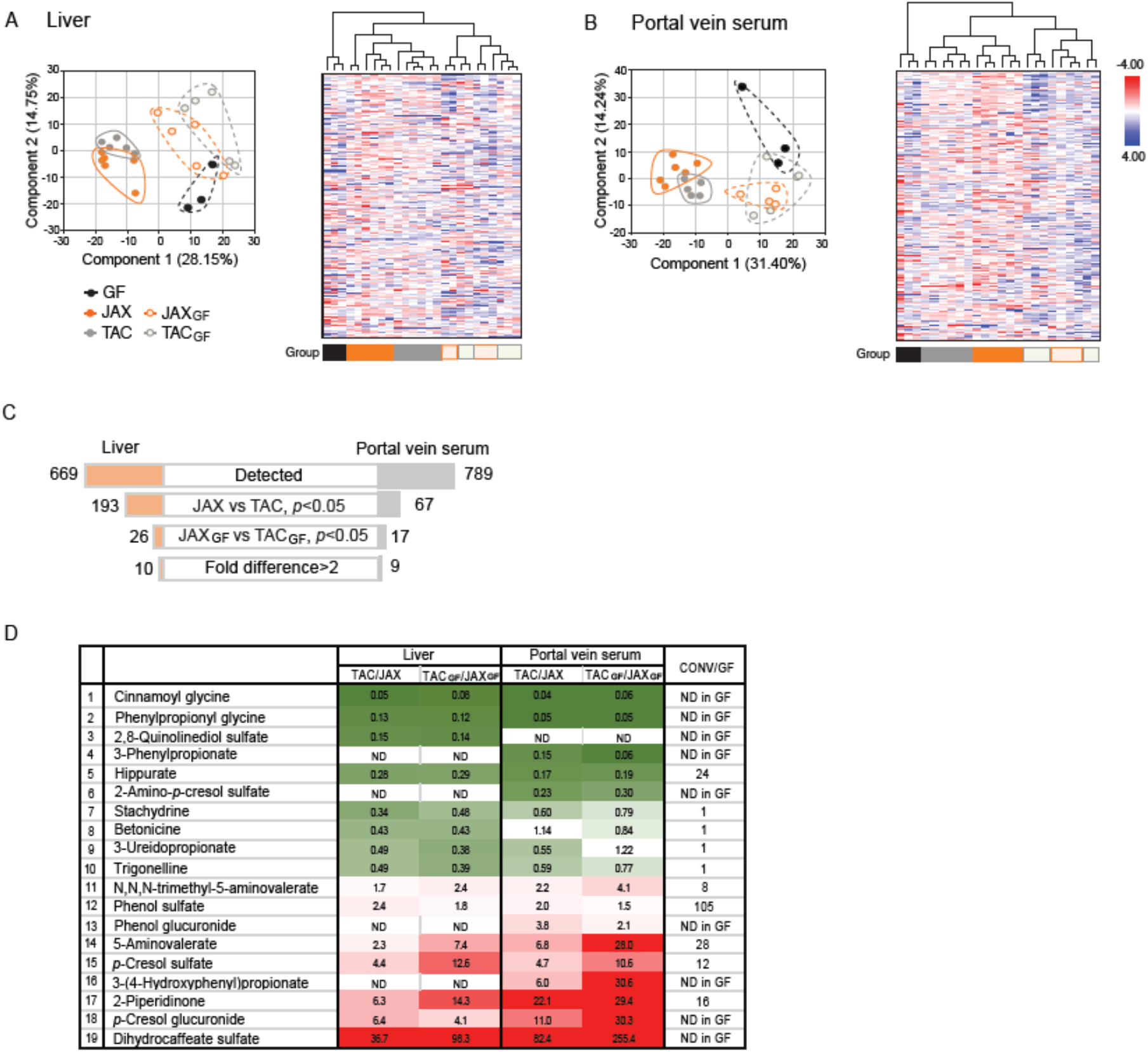
Certain gut microbe-derived metabolites are associated with altered susceptibility to APAP-induced hepatotoxicity. Liver and portal vein serum samples from JAX, TAC, GF, JAX_GF_, and TAC_GF_ were subjected to LC-MS/MS-based non-targeted metabolomic profiling. **A and B**. Principal component analysis and hierarchical clustering analysis of liver (A) and portal vein serum (B) are shown. **C**. Schematics of filtering steps to compile a list of metabolites in liver and portal vein serum samples that are associated with altered APAP hepatotoxicity. **D.** The signal ratios of metabolites that exhibited significant differences between JAX and TAC as well as between JAX_GF_, and TAC_GF_ mice are shown. CONV, conventional JAX or TC mice. ND, not detected.

To compile a list of gut microbiota-associated metabolites that are potentially linked to altered APAP hepatotoxicity, the metabolome data were filtered through multiple layers of selection criteria (Fig 3C). First, metabolites whose abundances differ between JAX and TAC (either liver or serum) samples were selected. They were further filtered by comparing their signals between JAX_GF_ and TAC_GF_ mice; only the metabolites that showed the same directional changes as in the conventional mice were selected. A total of 26 and 17 compounds, among which 10 and 9 metabolites showed >2-fold differences, were identified in the liver and portal vein serum samples, respectively. This led to compilation of a total 19 compounds as potential candidates for modulating APAP-induced hepatotoxicity (Fig 3D). Four compounds (i.e., stachydrine, betonicine, 3-ureidopropionate, and trigonelline) were readily detectable in the serum samples of GF mice, suggesting that host is a source. The rest were undetectable in GF serum or detected at less than 1/3 of those in conventional mice, suggesting that they may be produced by gut microbes. Consistent with results from the previous independent measurement (Fig S3A), the signals for *p*-cresol sulfate were higher in TAC than JAX mouse liver, supporting the robustness of the metabolome analysis.

Among 10 metabolites identified as being more abundant in the liver and portal vein serum of JAX mice, the following four were derived from the metabolic pathway of phenylalanine ^6^: 3-phenylpropionic acid (PPA), phenylpropionyl glycine, cinnamoyl glycine, and hippuric acid (Fig 4A). Phenylalanine is metabolized by gut bacteria to *trans*-cinnamic acid (tCA) and subsequently to PPA. Once absorbed, tCA and PPA are conjugated with glycine by hepatic glycine *N*-acyltransferase ^51^. PPA can also be further metabolized to benzoic acid, which is conjugated with glycine to form hippuric acid ^51^. To identify the sources for the greater abundance of these metabolites in JAX tissues, tCA and PPA contents in the mouse cecum were measured. tCA was undetectable in the cecum samples, likely due to the rapid conversion of tCA to PPA by gut microbiota ^6,52^. PPA levels, on the other hand, were readily detectable in the mouse cecum contents and comparable to those in feces from healthy individuals (i.e., 267 nmole/g feces) ^53^. PPA was undetected in GF mouse cecum contents, consistent with the idea that PPA is a gut microbial product. PPA levels in JAX and JAX_GF_ mouse cecum were 3.3- and 10-fold higher than TAC and TAC_GF_, respectively, suggesting that gut microbiota composition governs the PPA levels in mouse cecum contents (Fig 4B). Similar results were obtained for PPA levels in systemic circulation and portal vein (Fig 4B and 4C). PPA levels between cecum and portal (or systemic) blood exhibited excellent correlations (Fig S4A), suggesting that cecum PPA levels govern the host exposure to PPA.

**Fig 4.**
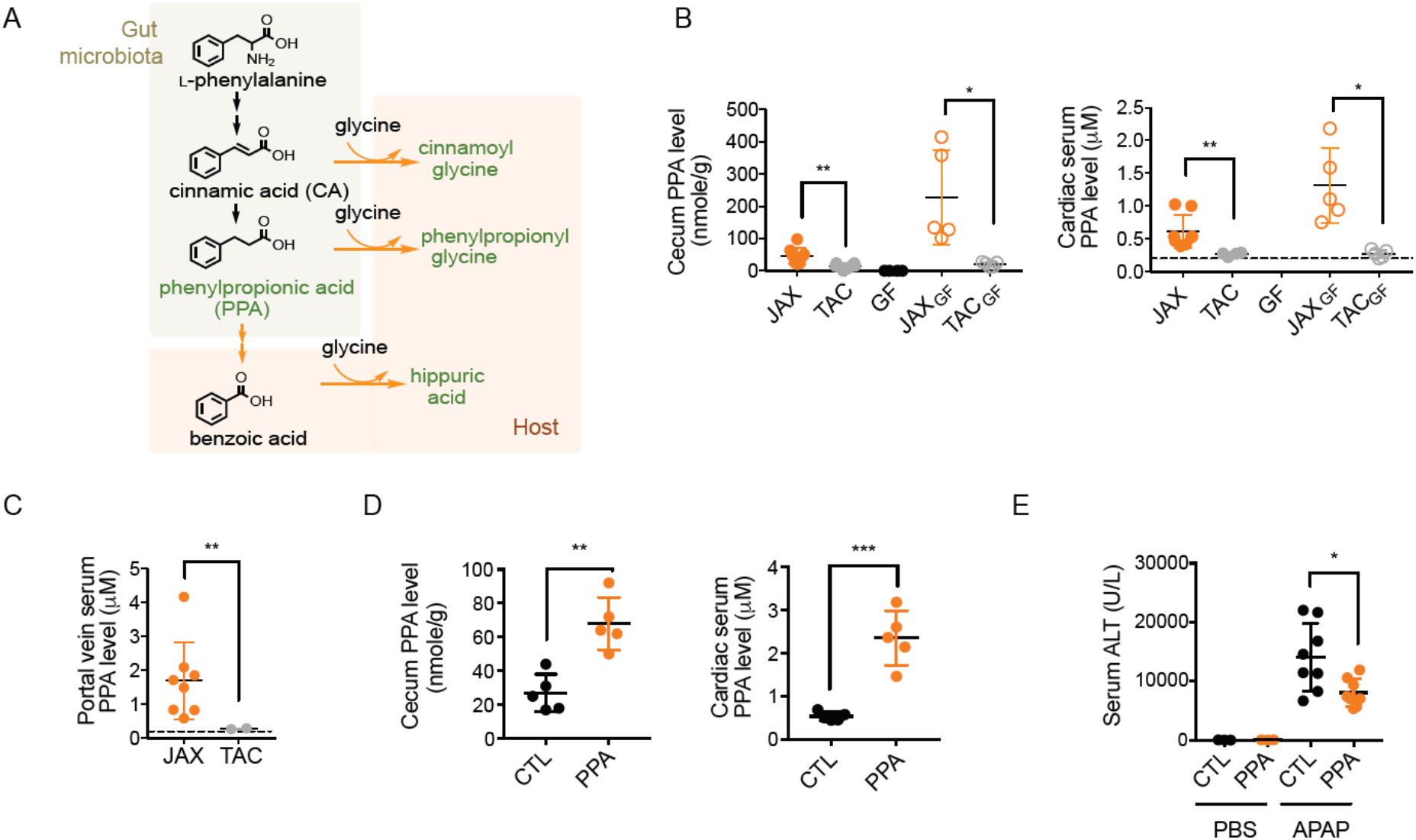
Gut microbe-associated metabolite, phenylpropionic acid (PPA), alleviates APAP hepatotoxicity. **A**. Gut microbe-mediated metabolic pathway of phenylalanine and host-mediated glycine conjugation. Green color, the metabolites identified to be associated with altered APAP toxicity. **B** and **C**. PPA levels in cecum contents and cardiac (B) or portal vein (C) serum of untreated mice were measured by LC-MS/MS. Dotted line denotes the limit of quantification (i.e., 0.2 μM). **D.** C57BL/6 mice from TAC were given access to PPA (0.4% in drinking water) or control (CTL) water for 4 weeks. PPA concentrations in cecum contents or serum prepared from cardiac blood were measured using LC-MS/MS. **E**. C57BL/6 mice from TAC were given access to PPA (0.4% in drinking water) or control (CTL) water for 4 weeks. The mice were then challenged with APAP (300 mg/kg or PBS, intraperitoneal injection) after overnight fasting, and sacrificed at 24 h post-dosing. All data are shown as mean ± S.D.

PPA exposure was greater in mice with JAX gut microbiota that is associated with lower APAP-induced hepatotoxicity as compared to mice with TAC microbiota. To determine whether PPA reduces the susceptibility to APAP hepatotoxicity, TAC mice were fed with PPA (0.4% in drinking water) for 4 weeks, followed by APAP administration. PPA amount in drinking water was determined based on (1) the average volume of daily water consumption for a mouse and (2) the previously reported pharmacokinetic information of tCA ^54^ (as relevant information for PPA is currently not available) to increase PPA exposure in TAC mice by about 5-fold. PPA supplementation had no effects on the amount of water intake or body weight gain of the mice (Fig S4B). The PPA dose used in this study led to ~3-fold increases in cecum and serum PPA levels in TAC mice (Fig 4D), the level approximating those in JAX mice. When the mice were challenged with APAP, the mice in PPA group exhibited significantly lower hepatotoxicity at 24 h post-dosing (Fig 4E), indicating that PPA alleviates APAP-induced hepatotoxicity.

### PPA represses basal hepatic CYP2E1 expression

To examine whether PPA supplementation pre-condition the liver towards lower APAP toxicity by modulating hepatic gene expression, transcriptomes of mouse liver tissues collected before APAP dosing were obtained using RNA-sequencing. Liver tissue samples collected at 6 h post-APAP dosing were also included to gain insights into hepatic responses to toxic doses of APAP. Results from PCA indicate that in both PPA and control groups, APAP treatment led to significant changes in hepatic gene expression (Fig S5A), in line with previous reports^55^. Importantly, basal gene expression profiles were similar between PPA and control groups, suggesting minimal effects of PPA supplementation on basal hepatic function. At 6 h post-dosing, however, hepatic expression of a large number of genes differed between PPA and control groups, potentially reflecting the biological consequences of reduced APAP hepatotoxicity by PPA.

CYP2E1 and CYP1A2 are major enzymes responsible for APAP conversion to NAPQI in mice ^56^. Basal protein expression levels of CYP2E1 (but not CYP1A2) in hepatic microsomes (the endoplasmic reticulum fraction) were about 2-fold lower in PPA group (Fig 5A). Consistent with the difference observed in CYP2E1 protein levels, APAP bioactivation (as measured by the formation rate of APAP-GSH upon incubation of hepatic microsome with APAP and GSH) was 1.5-fold lower in the PPA group (Fig 5B). In addition to endoplasmic reticulum, CYP2E1 proteins have been also detected in rat liver mitochondria ^57^; however, CYP2E1 proteins were undetectable in mitochondria fractions isolated from mouse liver tissues (Fig S5B). Of note, the basal hepatic mRNA levels of Cyp2e1 were similar between the control and PPA groups (Fig S5C), suggesting post-transcriptional regulation of CYP2E1 expression by PPA. Together, these results suggest that PPA alleviates APAP hepatotoxicity in part by decreasing hepatic CYP2E1 protein levels.

**Fig 5.**
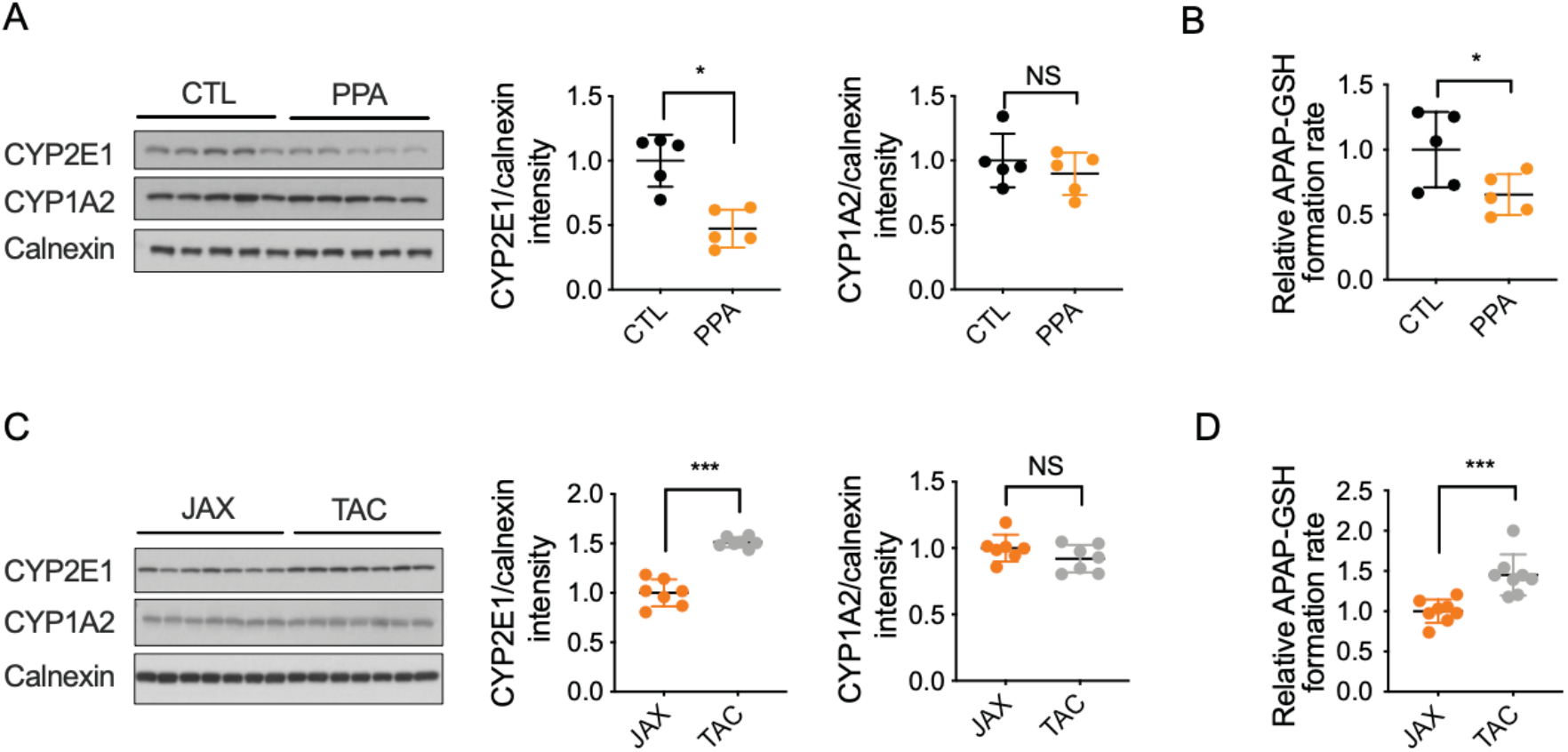
Phenylpropionic acid (PPA) alleviates APAP hepatotoxicity in part via decreased CYP2E1 expression. **A and B**. After PPA (or regular water as control, CTL) supplementation for 4 weeks, the mice were sacrificed after overnight fasting, and hepatic microsomes were prepared. (A) CYP2E1 and CYP1A2 protein levels in the microsomes were measured using western blot, using calnexin as a loading control. (B) Hepatic microsomes were incubated with APAP, and the extent of NAPQI formation was estimated by measuring APAP-GSH. **C and D.** C57BL/6 mice from JAX or TAC were sacrificed after overnight fasting. CYP2E1 protein expression in hepatic microsomes was measured using western blot. (D) The extent of NAPQI formation in hepatic microsomes was estimated by measuring APAP-GSH. All data are shown as mean ± S.D.

To determine the roles of PPA in altered susceptibility to APAP toxicity between mice with JAX or TAC gut microbiota, CYP2E1 protein levels in hepatic microsomes were compared between JAX and TAC mice as well as JAX_GF_ and TAC_GF_ mice. CYP2E1 protein levels were significantly lower in JAX or JAX_GF_ mice (Fig 5C and Fig S5D), with a corresponding decrease in APAP-GSH formation in hepatic microsomes from JAX mice (Fig 5D). The lower CYP2E1 activity in JAX as compared to TAC mice was also observed by using a CYP2E1 probe substrate, chlorzoxazone (Fig S5E). Previous studies have shown that degradation rates of CYP2E1 proteins are modulated by enzyme binding to substrates or inhibitors ^58–60^. PPA (up to 1 mM) did not inhibit CYP2E1-mediated chlorzoxazone hydroxylation in mouse hepatic microsomes while *p*-nitrophenol led to (Fig S5F), suggesting that PPA itself is unlikely a CYP2E1 substrate. These data support the notion that higher gut PPA levels and subsequent decreases in hepatic CYP2E1 expression may underlie reduced susceptibility to APAP hepatotoxicity in JAX mice.

## DISCUSSION

Gut microbiota produces thousands of chemically diverse small molecules ^2,3^, but the biological functions and the underlying molecular mechanisms for most are largely unknown. Here we report a novel function of a gut bacteria-derived metabolite PPA as a modulator of susceptibility to drug-induced liver injury in mice.

We found that C57BL/6 mice from TAC are more susceptible to APAP-induced hepatotoxicity than those from JAX. Similar differences in APAP toxicity between C57BL/6 mice from different vendors were previously reported ^61,62^ although the underlying mechanisms remained unclear. Genetic drifts between C57BL/6 mice from JAX and TAC have been reported ^44^, including the deletion mutation in nicotinamide nucleotide transhydrogenase (*Nnt)* in JAX mice. *Nnt* encodes an enzyme involved in mitochondrial antioxidant defense ^63^, a biological function linked to protection against APAP hepatotoxicity ^64^. In this study, however, JAX mice exhibited less hepatotoxicity after APAP administration. Furthermore, cohousing of JAX and TAC abrogated the differences in the susceptibility to APAP toxicity, and transplantation of gut microbiota to GF mice recapitulated the differences in APAP toxicity between the conventional JAX and TAC mice, indicating that susceptibility to APAP-induced hepatotoxicity is transferable by gut microbiota. These results strongly support the idea that differential gut microbiota contributes to the variations in the susceptibility to APAP-induced hepatotoxicity in mice.

Gut microbiota-derived metabolites mediate multiple aspects of gut microbe-host interactions. *p*-Cresol is a bacterial product of tyrosine that is readily conjugated by host sulfonation enzymes, and the product *p*-cresol sulfate is eliminated by urinary excretion ^65^. After a therapeutic dose of APAP was administered, *p*-cresol sulfate in urine was negatively correlated with urinary APAP-sulfate contents in humans ^23^, suggesting that *p*-cresol may compete with APAP for sulfonation. We found, however, that the hepatic levels of APAP-sulfate were 1000-fold higher than those of *p*-cresol sulfate and similar between JAX and TAC mice, indicating that the competition (if it occurs) is overcome when high toxic doses of APAP are administered. The contribution of *p*-cresol to altered susceptibility to APAP-induced hepatotoxicity observed in this study is minimal if any. Previous studies have shown that higher levels of 1-phenyl-1,2-propanedione (PPD) at ZT12 lead to enhanced APAP-induced hepatotoxicity in part by decreasing hepatic GSH reservoir in mice ^25^. PPD is a gut bacteria product ^25^ and also a volatile flavoring ingredient present in plants ^66^. Likely due to further metabolism by gut microbes ^25^, PPD levels in the liver and cecum contents were reported to be very low (~50 pmol/g cecum or liver) and below the detection limits in our assays. Potential involvement of PPD in altered susceptibility to APAP toxicity remains to be examined.

The non-targeted metabolomic analyses of liver and portal serum samples collected from the conventional and gut microbiota-transplanted mice revealed higher metabolic activities of phenylalanine in JAX gut microbiota. PPA levels in mouse cecum contents exhibited excellent positive correlations against those in the portal vein and systemic circulation, suggesting that PPA production by gut microbiota governs PPA exposure in the host. While the biological functions of PPA remain largely unknown, structural analogues of PPA have been reported to confer protection against chemical-induced liver injuries. 4-Phenylbutyric acid administered (30 min) after APAP dosing alleviated APAP-induced hepatotoxicity in mice, in part by reducing endoplasmic reticulum stress ^67^. tCA, the precursor of PPA, reversed ethanol-induced hepatotoxicity in mice by lowering hepatic expression of inflammatory mediators and repressing ethanol-mediated induction of CYP2E1 protein levels ^68^. PPA also decreases CYP2E1 protein (but not mRNA) levels, suggesting shared mechanisms between tCA and PPA in regulating CYP2E1 protein levels. Previous studies indicate that CYP2E1 substrates increases CYP2E1 protein levels via reducing CYP2E1 degradation mediated by ubiquitin-proteasome pathways ^58,59,69^. On the contrary, certain substrates such as carbon tetrachloride increase CYP2E1 degradation rate by forming reactive intermediates that can damage the protein ^70^. Our results indicate that PPA is not a CYP2E1 substrate or capable of inhibiting CYP2E1 enzyme activities but may decrease the levels of endogenous CYP2E1 substrates.

PPA can be produced via different routes that involve the interaction between diet and gut bacteria as well as the interaction between gut bacteria. First, a reductive pathway in the gut bacterium *Clostridium sporogenes* is shown to metabolize phenylalanine into PPA ^6,71^. The genes encoding enzymes mediating this metabolism are also present in other gut bacteria such as *Peptostreptococcus anaerobius* ^72^. Therefore, the amount of PPA produced by gut bacterial metabolism of phenylalanine will be dependent on total numbers of gut bacteria harboring the metabolic pathway genes and the availability of the substrate phenylalanine. Free phenylalanine in diet is absorbed mostly in the small intestine ^73^, and only a small fraction reaches the lower gastrointestinal tract. A large portion of phenylalanine available to gut bacteria in the large intestine is derived from proteolytic degradation of proteins by gut bacteria, which releases free phenylalanine or phenylalanine-containing peptides ^74^. Another source of phenylalanine in the large intestine is the biosynthesis of phenylalanine by gut bacteria ^75^. A recent study has shown that a major commensal gut bacteria, *Bacteroides thetaiotaomicron*, synthesizes *de novo* and excretes relatively large amounts of phenylalanine into the gut lumen, and it exhibits high strain variation in production capacity ^76^. Therefore, the availability of phenylalanine to PPA-producing gut bacteria will depend not only on diet but also the composition of neighboring gut bacteria.

Second, it has been known that certain plant-derived chemicals, flavonoids, are degraded by gut bacteria into PPA. For example, isoflavones such as apigenin and naringenin are shown to be metabolized into 3-(4-hydroxyphenyl)propionic acid (4-HPPA) by the gut bacterium *Clostridium orbiscindens* ^77^, and dehydroxylation of 4-HPPA by other gut bacteria results in formation of PPA ^78,79^. Apigenin, for example, exists as glycosides (e.g., apigenin-7-glucoside; A7G) in many vegetables, herbs, and beverages ^80^, and its availability to *C. orbiscindens* also requires prior removal of the sugar by gut bacterial glycosidases ^81^. Thus, the contribution of flavonoids such as apigenin to PPA production in the large intestine is governed by the interaction between the intake amounts of flavonoid glycosides, gut bacteria liberating the aglycone, and gut bacteria metabolizing the aglycone.

In conclusion, we have identified the gut bacterial metabolite PPA as one determinant of susceptibility to APAP-induced hepatotoxicity. Systemic levels of PPA appear to be well correlated with the amounts of PPA produced in the gut, perhaps suggesting the possibility of manipulating gut microbiota for PPA production.

## ACKNOWLEDGMENTS

This work was funded by the Chicago Biomedical Consortium with support from the Searle Funds at The Chicago Community Trust.

---

**Fig S1.**
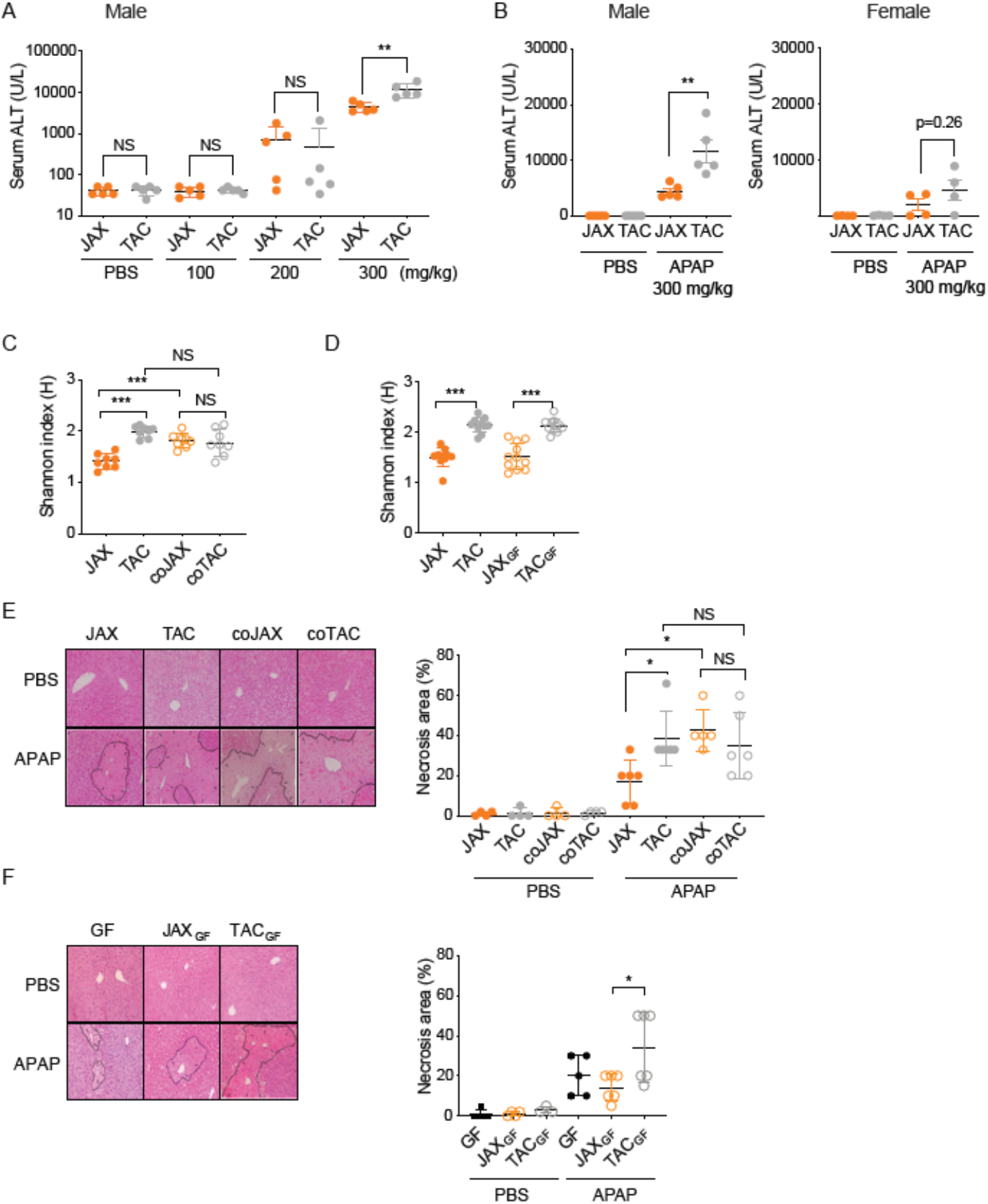
**A**. Male C57BL/6 mice from JAX or TAC received APAP (100, 200 or 300 mg/kg) or PBS intraperitoneally after overnight fasting. Mice were sacrificed after 24 h, and serum ALT levels were measured as a measure of hepatotoxicity. **B**. Male or female C57BL/6 mice received APAP (300 mg/kg or PBS, intraperitoneal injection) after overnight fasting, and serum ALT levels at 24 h post-dosing were measured. **C**. Gut microbiome was analyzed by 16S rRNA gene amplicon sequencing using fecal samples collected at the end of cohousing (4-week) or gut microbiota transplantation (oral gavage, followed by 4-week conventionalization). **E**. At the end of cohousing or gut microbiota transplantation, APAP (300 mg/kg or PBS, intraperitoneal injection) was administered after overnight fasting, and mice sacrificed after 24 h. Liver tissue was H/E stained.

**Fig S2.**
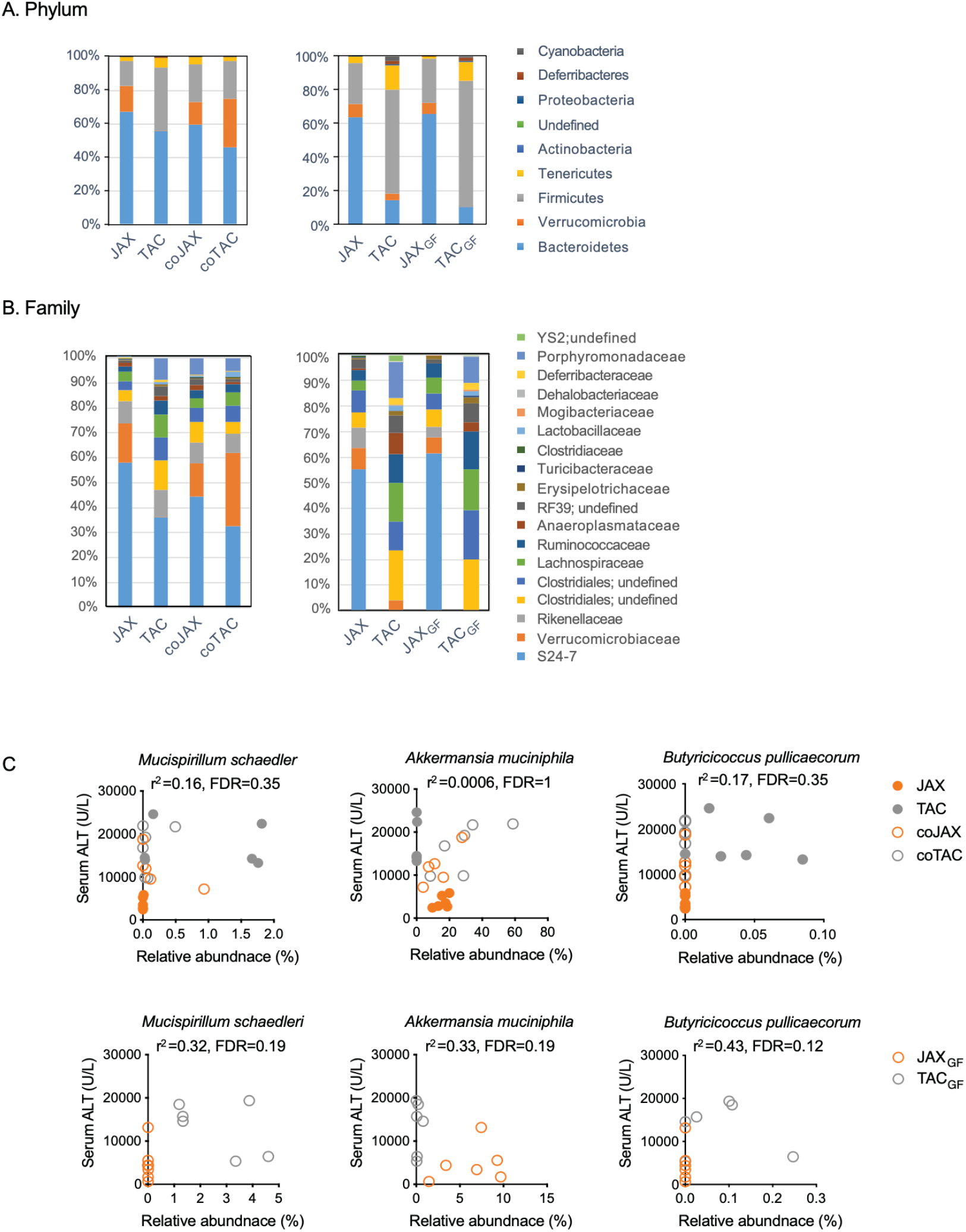
**A and B.** Gut microbiome of mice from co-housing (left) or gut microbiota-transplantation (right) experiments was analyzed by 16S rRNA gene amplicon sequencing. Gut bacteria compositions at phylum (A) and family (B) levels are shown. Bacteria taxa with less than 0.1% abundance are not shown. **C.** Correlations between fecal abundances of *M. schaedleri* or *A. muciniphila* and serum ALT levels in co-housed or gut microbiota-transplanted mice are shown.

**Fig S3.**
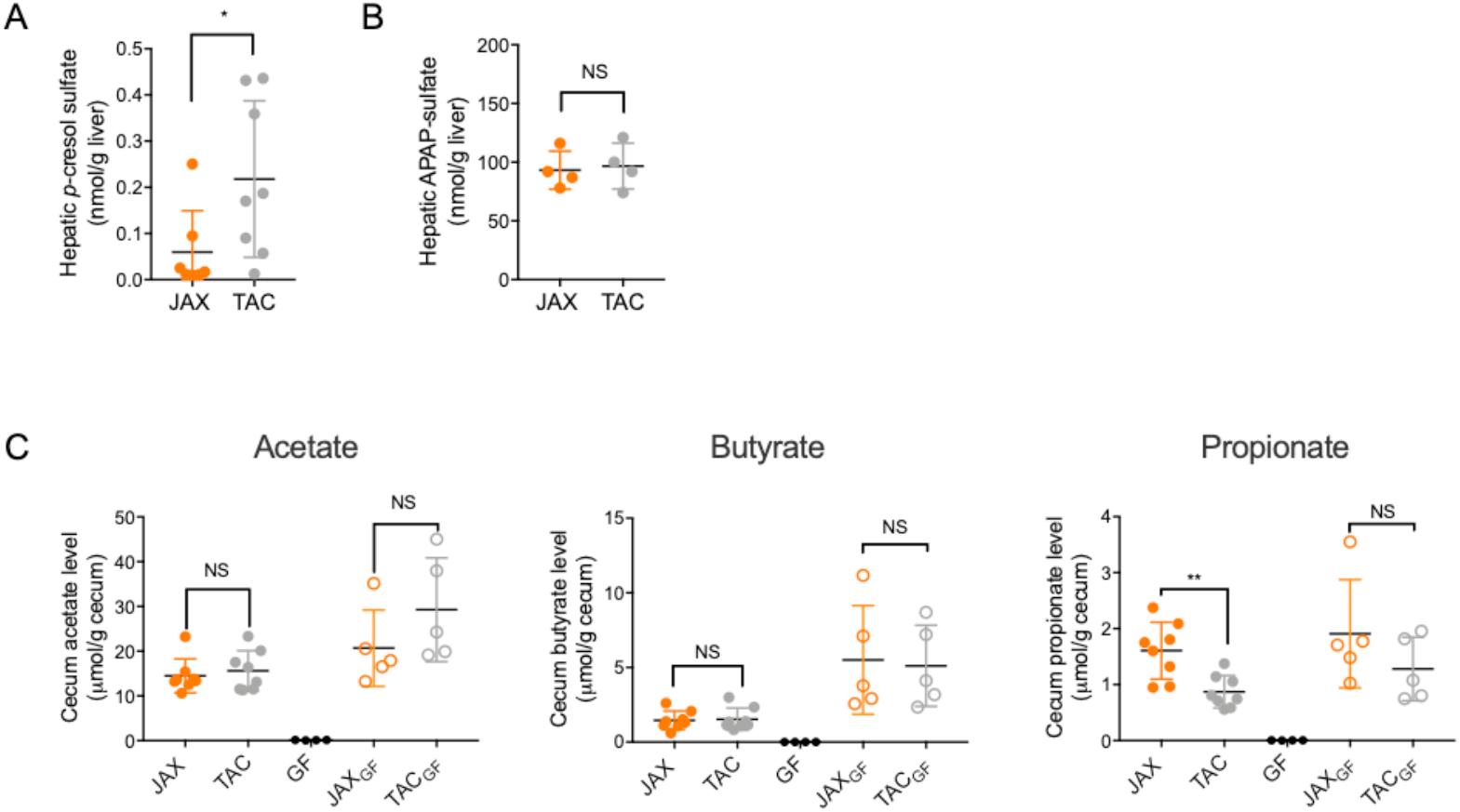
**A.** JAX or TAC mice were sacrificed after overnight fasting and liver tissues were collected. *p*-Cresol sulfate levels were measured using LC-MS/MS. **B.** APAP (300 mg/kg, intraperitoneal injection) was administered to C57BL/6 mice from JAX or TAC, and the mice were sacrificed after 15 min. APAP-sulfate levels in the liver were measured using LC-MS/MS. **C**. Conventional JAX or TAC mice as well as gut microbiota-transplanted mice were sacrificed after overnight fasting and cecum contents were collected. The levels of major short chain fatty acids in the cecum contents were measured using LC-MS/MS.

**Fig S4.**
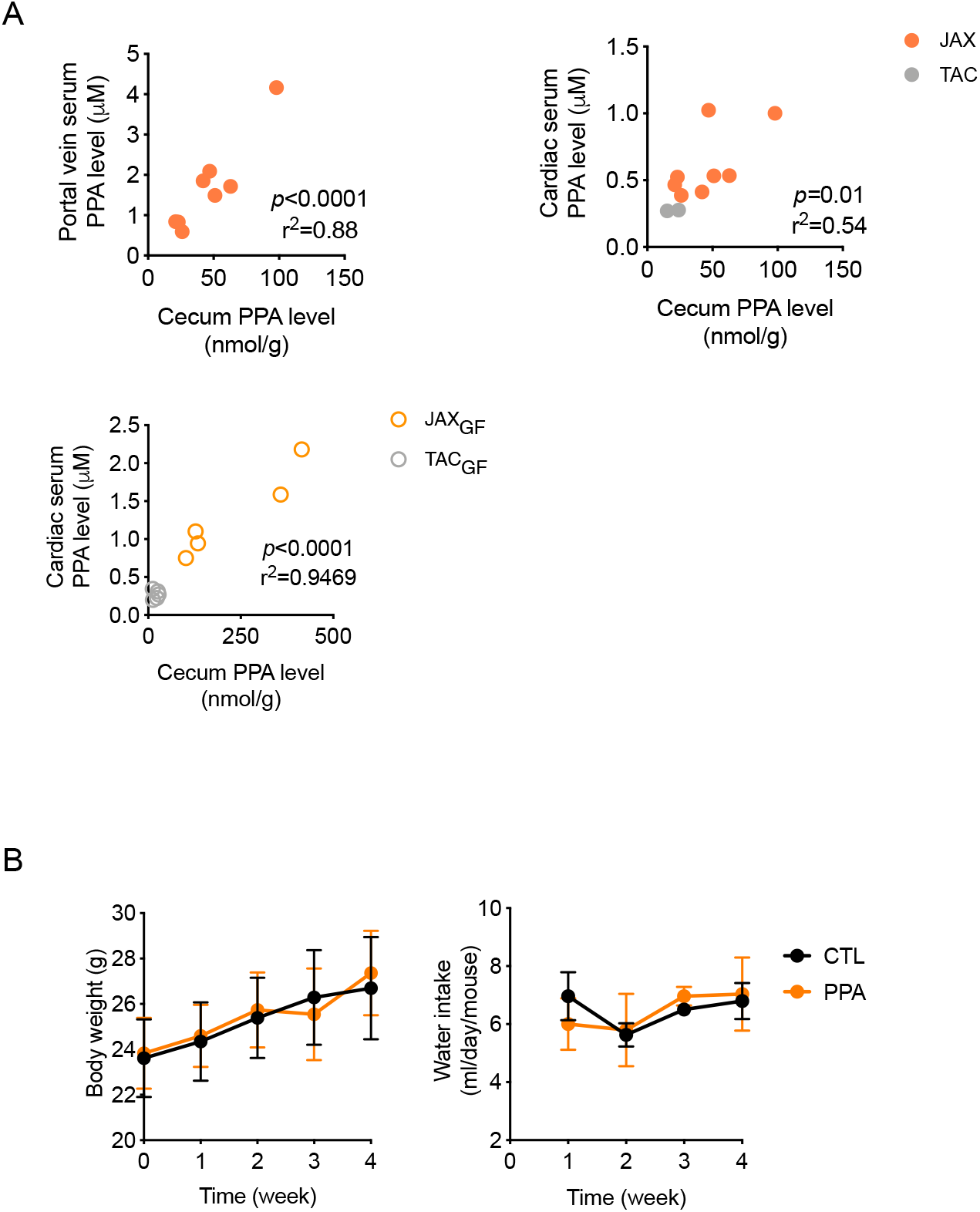
**A.** Correlations between PPA concentrations in cecum contents and serum prepared from portal vein (left) and cardiac blood (right) of JAX and TAC mice are shown. (bottom) Correlation between PPA concentrations in cecum contents and cardiac serum of JAX_GF_ and TAC_GF_ mice is shown. Mice that exhibited PPA concentrations below the quantification limit (i.e., 0.2 μM) are not included. **B**. TAC mice were fed with PPA (0.4% in drinking water) or regular water (CTL) for 4 weeks. Average body weight gain and water intake volume over time are shown. All data are shown as mean ± S.D.

**Fig S5.**
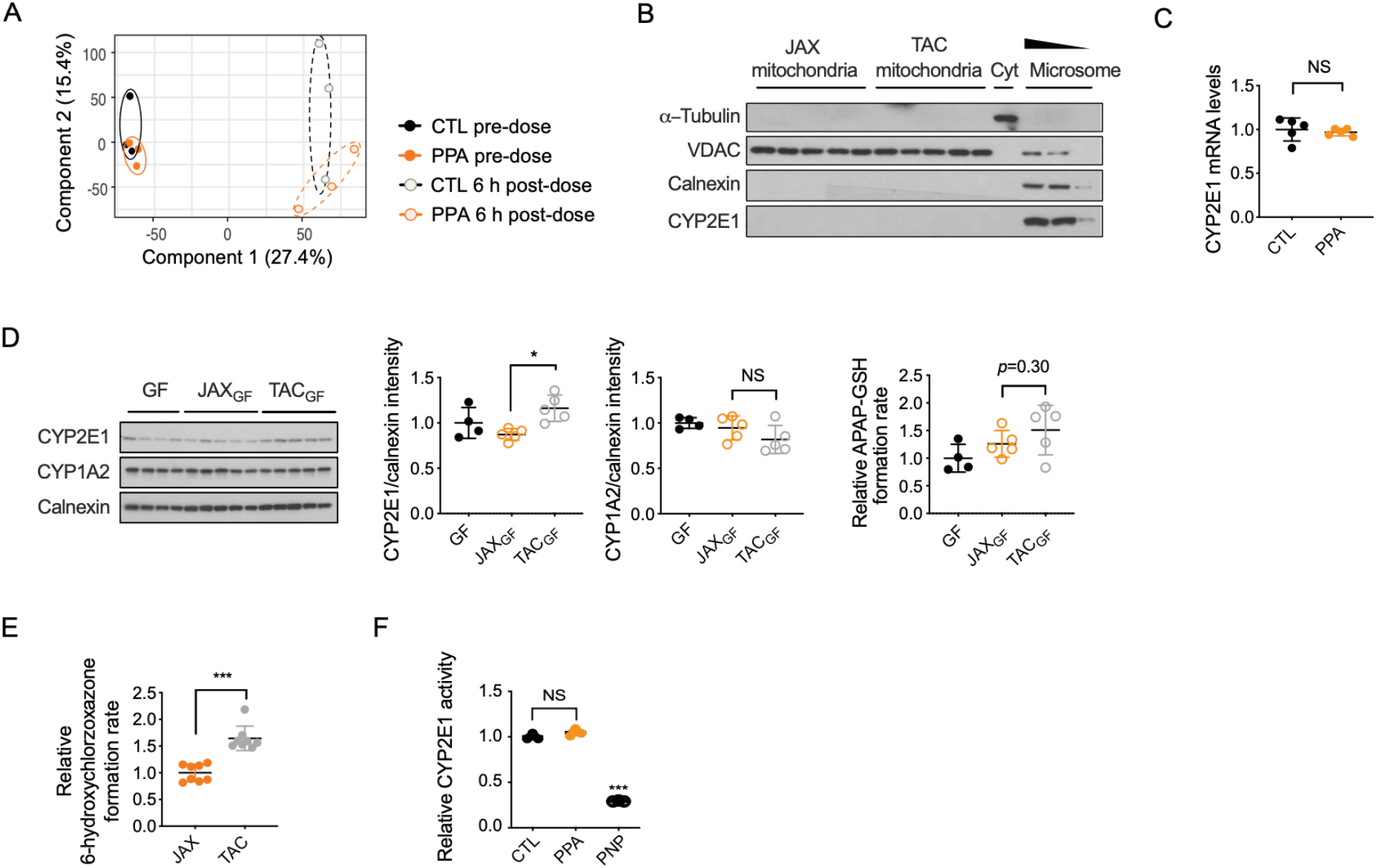
**A.** TAC mice were fed with PPA (0.4% in drinking water) or regular water (CTL) for 4 weeks. A group of mice were sacrificed after overnight fasting, and liver RNAs were isolated. The other group of mice received APAP (300 mg/kg intraperitoneally) after overnight fasting. The mice were sacrificed after 6 h, followed by liver RNA isolation. Transcriptome profiles were obtained by RNA-sequencing TAC mice. **B.** JAX or TAC mice were sacrificed after overnight fasting, followed by isolation of mitochondria, cytosol (Cyt) and microsomes. Mitochondrial and cytosolic proteins (10 μg/well) as well as microsomal proteins (1, 5, or 10 μg/well) were analyzed by western blot using antibodies detecting VDAC (voltage-dependent anion channel; mitochondria marker), α-tubulin (cytosol marker), and calnexin (microsome marker). **C.** Basal hepatic Cyp2e1 mRNA levels in PPA-supplemented mice were measured by qRT-PCR. **D.** Gut-microbiota transplanted mice were sacrificed after overnight fasting, and microsomal CYP2E1 and CYP1A2 levels were measured by western blot (left). APAP bioactivation levels (right) were determined by measuring APAP-GSH formation rate in the hepatic microsomes. **E.** Hepatic CYP2E1 activity levels were determined by measuring the rate of chlorzoxazone to 6-hydroxychlorzoxazone conversion rate in the hepatic microsomes. **F.** Mouse hepatic microsomes were incubated with chlorzoxazone (100 μM) in the presence of PPA (1 mM) or *p*-nitrophenol (PNP; 1 mM; a known CYP2E1 substrate as a competitive inhibitor) for 30 min, and the extent of 6-hydroxychlorzoxazone production was compared. All data are shown as mean ± S.D.

## REFERENCES

1. Qin, J., et al. A human gut microbial gene catalogue established by metagenomic sequencing. Nature 464, 59–65 (2010).

2. Donia, M.S. & Fischbach, M.A. HUMAN MICROBIOTA. Small molecules from the human microbiota. Science 349, 1254766 (2015).

3. Marcobal, A., et al. A metabolomic view of how the human gut microbiota impacts the host metabolome using humanized and gnotobiotic mice. The ISME journal 7, 1933–1943 (2013).

4. Cummings, J.H., Pomare, E.W., Branch, W.J., Naylor, C.P. & Macfarlane, G.T. Short chain fatty acids in human large intestine, portal, hepatic and venous blood. Gut 28, 1221–1227 (1987).

5. Husted, A.S., Trauelsen, M., Rudenko, O., Hjorth, S.A. & Schwartz, T.W. GPCR-Mediated Signaling of Metabolites. Cell Metab 25, 777–796 (2017).

6. Dodd, D., et al. A gut bacterial pathway metabolizes aromatic amino acids into nine circulating metabolites. Nature 551, 648–652 (2017).

7. Venkatesh, M., et al. Symbiotic bacterial metabolites regulate gastrointestinal barrier function via the xenobiotic sensor PXR and Toll-like receptor 4. Immunity 41, 296–310 (2014).

8. Holmes, E., et al. Human metabolic phenotype diversity and its association with diet and blood pressure. Nature 453, 396–400 (2008).

9. Liu, R., et al. Gut microbiome and serum metabolome alterations in obesity and after weight-loss intervention. Nat Med 23, 859–868 (2017).

10. Roberts, A.B., et al. Development of a gut microbe-targeted nonlethal therapeutic to inhibit thrombosis potential. Nat Med 24, 1407–1417 (2018).

11. Wallace, B.D., et al. Alleviating cancer drug toxicity by inhibiting a bacterial enzyme. Science 330, 831–835 (2010).

12. Isabella, V.M., et al. Development of a synthetic live bacterial therapeutic for the human metabolic disease phenylketonuria. Nat Biotechnol 36, 857–864 (2018).

13. Fischbach, M.A. Microbiome: Focus on Causation and Mechanism. Cell 174, 785–790 (2018).

14. Larson, A.M., et al. Acetaminophen-induced acute liver failure: results of a United States multicenter, prospective study. Hepatology 42, 1364–1372 (2005).

15. Lee, W.M. Acute liver failure in the United States. Semin Liver Dis 23, 217–226 (2003).

16. Ramachandran, A. & Jaeschke, H. Acetaminophen Hepatotoxicity. Semin Liver Dis 39, 221–234 (2019).

17. Kaplowitz, N. Acetaminophen hepatoxicity: what do we know, what don’t we know, and what do we do next? Hepatology 40, 23–26 (2004).

18. Watkins, P.B., et al. Aminotransferase elevations in healthy adults receiving 4 grams of acetaminophen daily: a randomized controlled trial. Jama 296, 87–93 (2006).

19. Schmidt, L.E., Dalhoff, K. & Poulsen, H.E. Acute versus chronic alcohol consumption in acetaminophen-induced hepatotoxicity. Hepatology 35, 876–882 (2002).

20. Whitcomb, D.C. & Block, G.D. Association of acetaminophen hepatotoxicity with fasting and ethanol use. Jama 272, 1845–1850 (1994).

21. Navarro, S.L., et al. UGT1A6 and UGT2B15 polymorphisms and acetaminophen conjugation in response to a randomized, controlled diet of select fruits and vegetables. Drug Metab Dispos 39, 1650–1657 (2011).

22. Court, M.H., et al. Candidate gene polymorphisms in patients with acetaminophen-induced acute liver failure. Drug Metab Dispos 42, 28–32 (2014).

23. Clayton, T.A., Baker, D., Lindon, J.C., Everett, J.R. & Nicholson, J.K. Pharmacometabonomic identification of a significant host-microbiome metabolic interaction affecting human drug metabolism. Proceedings of the National Academy of Sciences of the United States of America 106, 14728–14733 (2009).

24. Thaiss, C.A., et al. Microbiota Diurnal Rhythmicity Programs Host Transcriptome Oscillations. Cell 167, 1495–1510.e1412 (2016).

25. Gong, S., et al. Gut microbiota mediates diurnal variation of acetaminophen induced acute liver injury in mice. Journal of hepatology 69, 51–59 (2018).

26. Sharma, S., Chaturvedi, J., Chaudhari, B.P., Singh, R.L. & Kakkar, P. Probiotic Enterococcus lactis IITRHR1 protects against acetaminophen-induced hepatotoxicity. Nutrition 28, 173–181 (2012).

27. Caporaso, J.G., et al. Ultra-high-throughput microbial community analysis on the Illumina HiSeq and MiSeq platforms. ISME J 6, 1621–1624 (2012).

28. Caporaso, J.G., et al. Global patterns of 16S rRNA diversity at a depth of millions of sequences per sample. Proceedings of the National Academy of Sciences of the United States of America 108 Suppl 1, 4516–4522 (2011).

29. Bolyen, E., et al. QIIME 2: Reproducible, interactive, scalable, and extensible microbiome data science. PeerJ Preprints 6, e27295v27291 (2018).

30. Callahan, B.J., et al. DADA2: high-resolution sample inference from Illumina amplicon data. Nature methods 13, 581 (2016).

31. Bokulich, N.A., et al. Optimizing taxonomic classification of marker-gene amplicon sequences with QIIME 2’s q2-feature-classifier plugin. Microbiome 6, 90 (2018).

32. Thakare, R., Chhonker, Y.S., Gautam, N., Alamoudi, J.A. & Alnouti, Y. Quantitative analysis of endogenous compounds. J Pharm Biomed Anal 128, 426–437 (2016).

33. Wieckowski, M.R., Giorgi, C., Lebiedzinska, M., Duszynski, J. & Pinton, P. Isolation of mitochondria-associated membranes and mitochondria from animal tissues and cells. Nat Protoc 4, 1582–1590 (2009).

34. Laine, J.E., Auriola, S., Pasanen, M. & Juvonen, R.O. Acetaminophen bioactivation by human cytochrome P450 enzymes and animal microsomes. Xenobiotica; the fate of foreign compounds in biological systems 39, 11–21 (2009).

35. Patten, C.J., et al. Cytochrome P450 enzymes involved in acetaminophen activation by rat and human liver microsomes and their kinetics. Chemical research in toxicology 6, 511–518 (1993).

36. McGill, M.R., et al. Plasma and liver acetaminophen-protein adduct levels in mice after acetaminophen treatment: dose-response, mechanisms, and clinical implications. Toxicology and applied pharmacology 269, 240–249 (2013).

37. McGill, M.R., et al. HepaRG cells: a human model to study mechanisms of acetaminophen hepatotoxicity. Hepatology 53, 974–982 (2011).

38. McCarthy, D.J., Chen, Y. & Smyth, G.K. Differential expression analysis of multifactor RNA-Seq experiments with respect to biological variation. Nucleic Acids Res 40, 4288–4297 (2012).

39. Robinson, M.D., McCarthy, D.J. & Smyth, G.K. edgeR: a Bioconductor package for differential expression analysis of digital gene expression data. Bioinformatics 26, 139–140 (2010).

40. Benjamini, Y. & Hochberg, Y. Controlling the False Discovery Rate - a Practical and Powerful Approach to Multiple Testing. J R Stat Soc B 57, 289–300 (1995).

41. McIntosh, C.M., Chen, L., Shaiber, A., Eren, A.M. & Alegre, M.-L. Gut microbes contribute to variation in solid organ transplant outcomes in mice. Microbiome 6(2018).

42. Sivan, A., et al. Commensal Bifidobacterium promotes antitumor immunity and facilitates anti-PD-L1 efficacy. Science (2015).

43. Du, K., Williams, C.D., McGill, M.R. & Jaeschke, H. Lower susceptibility of female mice to acetaminophen hepatotoxicity: Role of mitochondrial glutathione, oxidant stress and c-jun N-terminal kinase. Toxicol Appl Pharmacol 281, 58–66 (2014).

44. Almodovar, A.J., Luther, R.J., Stonebrook, C.L. & Wood, P.A. Genomic structure and genetic drift in C57BL/6 congenic metabolic mutant mice. Mol Genet Metab 110, 396–400 (2013).

45. McIntosh, C.M., Chen, L., Shaiber, A., Eren, A.M. & Alegre, M.L. Gut microbes contribute to variation in solid organ transplant outcomes in mice. Microbiome 6, 96 (2018).

46. Ivanov, II, et al. Induction of intestinal Th17 cells by segmented filamentous bacteria. Cell 139, 485–498 (2009).

47. Robertson, S.J., et al. Comparison of Co-housing and Littermate Methods for Microbiota Standardization in Mouse Models. Cell Rep 27, 1910–1919 e1912 (2019).

48. den Besten, G., et al. The role of short-chain fatty acids in the interplay between diet, gut microbiota, and host energy metabolism. J Lipid Res 54, 2325–2340 (2013).

49. Veech, R.L., et al. The effect of short chain fatty acid administration on hepatic glucose, phosphate, magnesium and calcium metabolism. Advances in experimental medicine and biology 194, 617–646 (1986).

50. Hara, H., Haga, S., Aoyama, Y. & Kiriyama, S. Short-chain fatty acids suppress cholesterol synthesis in rat liver and intestine. J Nutr 129, 942–948 (1999).

51. Badenhorst, C.P., Erasmus, E., van der Sluis, R., Nortje, C. & van Dijk, A.A. A new perspective on the importance of glycine conjugation in the metabolism of aromatic acids. Drug Metab Rev 46, 343–361 (2014).

52. Gutierrez-Diaz, I., et al. Could Fecal Phenylacetic and Phenylpropionic Acids Be Used as Indicators of Health Status? J Agric Food Chem 66, 10438–10446 (2018).

53. Colosimo, D.A., et al. Mapping Interactions of Microbial Metabolites with Human G-Protein-Coupled Receptors. Cell Host Microbe 26, 273–282 e277 (2019).

54. H.P., F., C.L., H., L.T., Z. & R.F., L. Pharmacokinetics and Bioavailability of Cinnamic Acid in Mice. J. China. Pharm. Univ. 35, 328–330 (2004).

55. Williams, D.P., et al. Time course toxicogenomic profiles in CD-1 mice after nontoxic and nonlethal hepatotoxic paracetamol administration. Chemical research in toxicology 17, 1551–1561 (2004).

56. Zaher, H., et al. Protection against acetaminophen toxicity in CYP1A2 and CYP2E1 double-null mice. Toxicology and applied pharmacology 152, 193–199 (1998).

57. Robin, M.A., et al. Mitochondrial targeted cytochrome P450 2E1 (P450 MT5) contains an intact N terminus and requires mitochondrial specific electron transfer proteins for activity. The Journal of biological chemistry 276, 24680–24689 (2001).

58. Song, B.J., Veech, R.L., Park, S.S., Gelboin, H.V. & Gonzalez, F.J. Induction of rat hepatic N-nitrosodimethylamine demethylase by acetone is due to protein stabilization. The Journal of biological chemistry 264, 3568–3572 (1989).

59. Roberts, B.J., Song, B.J., Soh, Y., Park, S.S. & Shoaf, S.E. Ethanol induces CYP2E1 by protein stabilization. Role of ubiquitin conjugation in the rapid degradation of CYP2E1. The Journal of biological chemistry 270, 29632–29635 (1995).

60. Pratt-Hyatt, M., Lin, H.L. & Hollenberg, P.F. Mechanism-based inactivation of human CYP2E1 by diethyldithocarbamate. Drug metabolism and disposition: the biological fate of chemicals 38, 2286–2292 (2010).

61. Bourdi, M., Davies, J.S. & Pohl, L.R. Mispairing C57BL/6 substrains of genetically engineered mice and wild-type controls can lead to confounding results as it did in studies of JNK2 in acetaminophen and concanavalin A liver injury. Chemical research in toxicology 24, 794–796 (2011).

62. Duan, L., et al. Differential susceptibility to acetaminophen-induced liver injury in sub-strains of C57BL/6 mice: 6N versus 6J. Food Chem Toxicol 98, 107–118 (2016).

63. Toye, A.A., et al. A genetic and physiological study of impaired glucose homeostasis control in C57BL/6J mice. Diabetologia 48, 675–686 (2005).

64. Ronchi, J.A., Francisco, A., Passos, L.A., Figueira, T.R. & Castilho, R.F. The Contribution of Nicotinamide Nucleotide Transhydrogenase to Peroxide Detoxification Is Dependent on the Respiratory State and Counterbalanced by Other Sources of NADPH in Liver Mitochondria. The Journal of biological chemistry 291, 20173–20187 (2016).

65. Smith, E.A. & Macfarlane, G.T. Enumeration of human colonic bacteria producing phenolic and indolic compounds: effects of pH, carbohydrate availability and retention time on dissimilatory aromatic amino acid metabolism. J Appl Bacteriol 81, 288–302 (1996).

66. Burdock, G.A. Encyclopedia of food and color additives, (CRC Press, Boca Raton, 1997).

67. Kusama, H., et al. Sodium 4-phenylbutyric acid prevents murine acetaminophen hepatotoxicity by minimizing endoplasmic reticulum stress. J Gastroenterol 52, 611–622 (2017).

68. Yan, S.L., Wang, Z.H., Yen, H.F., Lee, Y.J. & Yin, M.C. Reversal of ethanol-induced hepatotoxicity by cinnamic and syringic acids in mice. Food Chem Toxicol 98, 119–126 (2016).

69. Eliasson, E., Johansson, I. & Ingelman-Sundberg, M. Ligand-dependent maintenance of ethanol-inducible cytochrome P-450 in primary rat hepatocyte cell cultures. Biochemical and biophysical research communications 150, 436–443 (1988).

70. Tierney, D.J., Haas, A.L. & Koop, D.R. Degradation of cytochrome P450 2E1: selective loss after labilization of the enzyme. Archives of biochemistry and biophysics 293, 9–16 (1992).

71. Elsden, S.R., Hilton, M.G. & Waller, J.M. The end products of the metabolism of aromatic amino acids by Clostridia. Arch Microbiol 107, 283–288 (1976).

72. Wlodarska, M., et al. Indoleacrylic Acid Produced by Commensal Peptostreptococcus Species Suppresses Inflammation. Cell Host Microbe 22, 25–37 e26 (2017).

73. Broer, S. & Fairweather, S.J. Amino Acid Transport Across the Mammalian Intestine. Compr Physiol 9, 343–373 (2018).

74. Windey, K., De Preter, V. & Verbeke, K. Relevance of protein fermentation to gut health. Mol Nutr Food Res 56, 184–196 (2012).

75. Pittard, J. & Yang, J. Biosynthesis of the Aromatic Amino Acids. EcoSal Plus 3(2008).

76. Chen, H., et al. A Forward Chemical Genetic Screen Reveals Gut Microbiota Metabolites That Modulate Host Physiology. Cell 177, 1217–1231 e1218 (2019).

77. Schoefer, L., Mohan, R., Schwiertz, A., Braune, A. & Blaut, M. Anaerobic degradation of flavonoids by Clostridium orbiscindens. Appl Environ Microbiol 69, 5849–5854 (2003).

78. Mosele, J.I., Macia, A. & Motilva, M.J. Metabolic and Microbial Modulation of the Large Intestine Ecosystem by Non-Absorbed Diet Phenolic Compounds: A Review. Molecules 20, 17429–17468 (2015).

79. Braune, A. & Blaut, M. Bacterial species involved in the conversion of dietary flavonoids in the human gut. Gut Microbes 7, 216–234 (2016).

80. Bhagwat, S., Haytowitz, D.B. & Holden, J.M. USDA Database for the Flavonoid Content of Selected Foods. Release 3.1. in USDA Database for the Flavonoid Content of Selected Foods, Vol. 2019 (U.S. Department of Agriculture, Agricultural Research Service, 2014).

81. Hanske, L., Loh, G., Sczesny, S., Blaut, M. & Braune, A. The bioavailability of apigenin-7-glucoside is influenced by human intestinal microbiota in rats. J Nutr 139, 1095–1102 (2009).

